# Investigating Amino acid Enrichments and Patterns in Phase-Separating Proteins: Understanding Biases in Liquid-Liquid Phase Separation

**DOI:** 10.1101/2024.12.19.629394

**Authors:** Joana Calvário, Diogo Antunes, Rita Cipriano, Daniela Kalafatovic, Goran Mauša, Ana S. Pina

**Affiliations:** Instituto de Tecnologia Química e Biológica António Xavier, Universidade Nova de Lisboa, Av. da República, 2780-157 Oeiras, Portugal; University of Rijeka, Faculty of Biotechnology and Drug Development, 51000 Rijeka, Croatia; University of Rijeka, Center for Artificial Intelligence and Cybersecurity, 51000 Rijeka, Croatia; University of Rijeka, Faculty of Engineering, 51000, Rijeka, Croatia

**Keywords:** Membraneless Organelles, Liquid-Liquid Phase Separation (LLPS), Intrinsically Disordered Proteins (IDPs), Amino acid Enrichment, Motif discovery, Peptides, Minimalistic peptide design

## Abstract

Liquid-Liquid Phase Separation (LLPS) forms membraneless organelles, enhancing biochemical processes. The stickers-and-spacers model explains LLPS but is mainly validated in Prion-like RNA Binding Proteins. We explore peptide motifs in LLPS in broader protein contexts. We developed a computational approach for motif discovery, implemented in 178 Phase-Separating Proteins (PhSePs), complemented by the FuzDrop and CIDER servers, which identified droplet-promoting regions (DPRs) and examined disorder-related characteristics. Our database of PhSePs was analyzed against proteins with low propensity for LLPS. This comparative analysis revealed 129 enriched peptide motifs with folds higher than 0.2, consisting of 3 to 6 residues, with tetrapeptides being the most prevalent. Key features of the enriched motifs included Gly-rich sequences punctuated with aromatic, charged, and polar residues, as well as homopeptide repeats (e.g., GGDR, SRGG, YGGG, QQQQ, PPPP). Analysis of motif presence, frequency, and co-occurrence revealed widely distributed motifs across different DPRs, identified motifs with significant repetitive patterns, and highlighted motif trios that are more likely to co-occur within a sequence. By harvesting this analysis, we developed a data-driven approach for minimalistic peptide design with LLPS propensity, further using the CIDER server for peptide characterization and peptide design refinement. We designed 8 peptides with various motif combinations and amino acid distributions, which were experimentally validated to undergo LLPS, exhibiting liquid-like behavior with diverse molecular mobility patterns and droplet dynamics. Our approach bridges a non-biased computational approach with experimental validation, offering insights into sequence determinants of phase separation, with the potential for designing minimalistic synthetic condensates with tailored properties.

## Introduction

Compartmentalization is key in living systems, allowing for specialized environments via physical membranes or liquid-liquid phase separation (LLPS) and leading to the formation of membraneless organelles like nucleoli and stress granules. Such partitioning in cells enhances essential biochemical processes. ^1–5^ LLPS-derived organelles often comprise intrinsically disordered proteins/regions (IDPs/IDRs), known for their low-complexity amino acid regions (LCRs) and lack of a defined 3D structure, and nucleic acids. ^1–3,6^ The sequence composition and distribution of LCRs in IDPs significantly influences phase transitions, prompting recent research into how primary sequences affect LLPS. ^3,4,7^

The stickers and spacers model, inspired by associative polymers, explains LLPS in IDPs. “Stickers”, often aromatic amino acids (e.g., Tyr, Phe), interact via non-covalent bonds (hydrophobic, electrostatic, pi–pi, cation–pi, and hydrogen bonds), while “Spacers” are typically polar residues (e.g., Gly, Ser, Gln) that provide flexibility and a dynamic nature to the protein. ^1–3,5^ Consistent distribution of sticker residues and charged amino acid clusters promotes LLPS, while its absence may lead to amorphous precipitates, highlighting the critical role of multivalency, patterning, and charge distribution in driving protein phase separation. ^2,5,8^ The stickers and spacers model has proven valuable in understanding LLPS, yet its validation has primarily focused on prion-like RNA binding proteins, particularly the FET family (FUS, EWSR1, and TAF15). ^2,4,5,9^ This narrow focus on RNA binding proteins has limited our understanding of how protein context and function influence LLPS across diverse families, leaving uncertainty about whether such principles can be generalized to other cellular functions. The use of peptide-based model systems in synthetic environments has also been used to study of LLPS in a simplified manner, although continuing to focus on the stickers and spacers conceptual model. ^10–17^

Specific motifs related to LLPS have been identified, primarily through a combination of coincidental observations, targeted studies of known proteins, and bioinformatic analysis of existing protein databases, including GAR (Glycine- and arginine-rich), YGG motifs, and proline-rich regions. ^18–20^

Recent LLPS studies employ specialized computational tools. Predictors ^21,22^ use machine learning to identify phase-separating proteins, web servers ^23–25^ analyze physicochemical properties, while databases ^26–28^ catalog verified LLPS proteins. These resources aid in identifying and characterizing LLPS-prone proteins. Despite significant advancements in artificial intelligence and algorithmic approaches for studying LLPS, the sequence-LLPS relationship in a context-dependent manner is still overlooked. Current algorithms often exhibit biases towards specific protein families or motifs, which constrains their predictive capabilities. Furthermore, existing approaches have limitations in their ability to rationally design novel LLPS-prone sequences, as these have been primarily focused on the stickers and spacers model.

In this work, we aim to contribute for the understanding of LLPS rules across diverse protein classes, considering both functionality and context, and further design synthetic minimalistic models based on peptide systems. We have developed a computational approach for the discovery of peptide motifs by using a database comprising of 178 Phase-Separating Proteins (PhSePs) categorized by function. Our approach was complemented by two widely recognized servers in the field of disordered proteins and LLPS: the FuzDrop server, introduced by Fuxreiter and Vendruscolo, ^24,25^ and the CIDER (Classification of Intrinsically Disordered Ensemble Regions) server, developed by the Pappu Lab. ^23,29,30^ The FuzDrop server is an invaluable resource for elucidating the principles of phase transitions and likelihood of liquid-like droplet formation through its predictive analysis of local sequence properties such as composition, hydrophobicity, and structural disorder. We utilized FuzDrop to predict the probability of proteins undergoing LLPS and to pinpoint Droplet Promoting Regions (DPRs) within their sequences. The CIDER server is an essential tool for analyzing intrinsic disorder-related characteristics, and has been used to study the distribution of such parameters across all IDRs in the human proteome, providing insights into their potential phase separation behavior. ^31^ We used CIDER to examine several parameters in DPRs, including charge distribution, hydropathy, and fraction of disorder-promoting residues, while also comparing these metrics with the NODPR regions of PhSePs.

Our approach encompassed the analysis of amino acid composition and their properties in PhSePs and DPRs, as well as the identification and evaluation of statistically significant patterns of peptide motifs (ranging from 3 to 6 residues in length) through comparison with the same peptide motifs in a negative control database containing proteins that do not phase separate.

The presence (number of distinct DPRs with at least one instance of a peptide motif), frequency (total incidences of a peptide motif across all DPRs, including repeats within sequences), and co-occurrence (number of DPRs where three peptide motifs coexist) of the peptide motifs were evaluated. While the presence and frequency provide insights about the importance of peptide motif distribution within a sequence, the co-occurrence of the motifs provide the strategy to design synthetic peptides composed of 10 to 14 residues, by optimizing the sequence space of the designed minimalistic peptides. This design allowed us to incorporate various motif combinations and amino acid distributions, for subsequent experimental validation.

## Materials and Methods

### 1. Deciphering Amino Acid Motif Patterns in Phase-Separating Proteins

For the present study we employed two databases, comprising a total of 178 distinct PhSePs sourced from LLPSDB ^26,27^ and PhaSePro ^28^. The selection of these databases was driven by their robust experimental background and thorough validation of phase separation behaviors. The Phase-Separating Proteins (PhSePs) were classified based on their functionality, including RNA binding, DNA binding, Chromatin binding, Regulation, Hydrolase, and Structure proteins, with respective counts of 79, 42, 21, 27, 17, 10 proteins.

We used the FuzDrop server ^24,25^ to identify Droplet Promoting Regions (DPRs) and regions that do not promote droplet formation (NODPRs). The resulting 712 DPRs in our database were, on average, 72.1 amino acids in length, while the 702 NODPRs had an average of 90.2 residues.

To ensure statistical reliability of our subsequent analysis, we created a negative control dataset. This dataset was curated from the Universal Protein Resource (UniProt) and consisted initially of 3,000 human proteins. Each protein in this dataset has a length ranging from 400 to 800 residues, comparable to our primary dataset of PhSePs, which have an average length of around 600 residues. We processed these 3,000 proteins through the FuzDrop server. Subsequently, we randomly selected 208 proteins that demonstrated a droplet-promoting probability (pDP) below 20%, referred as non-PhSePs. On average, the 208 proteins from the negative database exhibited 0 to 4 DPRs, with a mean value of 0.92 DPRs per protein.

For the following computational analysis, we used Python 3.11 running on Spyder 5.4.3 from Anaconda navigator 2.5.2. To quantify the composition of amino acid residues and their character, we created the “STATITIAN” script, which calculates the frequencies of all 20 amino acids within the full protein sequences and separately within DPRs and NODPRs. This analysis provided both count and percentage distributions of the amino acids, along with a detailed account of the side chain properties for each residue. The same method was additionally applied to each protein family as well as non-PhSePs.

For further analysis of our database, we employed the CIDER (Classification of Intrinsically Disordered Ensemble Regions) algorithm, specifically using localCIDER Version 0.1.18. This analysis focused on five key parameters, the Fraction of Charged Residues (FCR), Net Charge per Residue (NCPR), Kappa (κ), Mean Hydropathy and the Fraction of Disorder Promoting Residues. The program utilized in this step, “CIDER FOR PROTEINS”, is available in our GitHub repository.

To study amino acid patterning in PhSePs, we conducted a thorough analysis of all protein sequences, focusing on DPR regions. We developed the “SELECTOR” script to analyze DPR sequences for all possible motifs of a specified length. By providing an input of the target peptide motif length (in our study we used a length of 3 to 6 residues for a minimalistic approach), our program identifies all contiguous subsequences of the specified length within the DPR database. The “SELECTOR” script simultaneously computes a motifs’ presence (number of distinct DPRs with at least one instance of a motif) and frequency (total occurrences across all DPRs, including repeats within sequences). This program was used in our research for motif discovery in DPRs of PhSePs, the proteins by family, as well as non-PhSePs. Fold values were calculated by dividing the presence/frequency of a given motif in the DPR sequences of PhSePs for the same parameters of the negative database of non-PhSePs (i.e. Motif presence = Presence in DPRs of PhSePs/Presence in non-PhSePs, and Motif frequency = Frequency in DPRs of PhSePs/frequency in non-PhSePs). The resulting fold values were then normalized on a scale of 0 to 1, thus returning a presence fold (PF) and a frequency fold (FF). Recognizing that both the presence and frequency of motifs are crucial for understanding the mechanism of LLPS, we assigned them equal importance, by calculating a combined fold score (CF), where CF = 0.5 PF + 0.5 FF. To identify the most significant and frequently occurring motifs, we focused on motifs with a CF of 0.2 or higher. As a result, 129 motifs met this condition. The presence and frequency of the 129 discovered motifs were additionally computed in NODPRs sequences to validate their enrichment in DPRs.

We developed the “FREQUENCY” script which enabled us to further assess presence and frequency values per individual DPR sequences, both in PhSePs and in protein families.

All the described scripts can be found in our GitHub repository.

### 2. Minimalistic peptide design based on motif co-occurrence

To design minimalistic peptides with LLPS propensity based on the co-occurrence of motif trios, we developed the “COMBINER” script, which analyzes combinations of the 129 discovered peptide motifs. Our analysis specifically focused on combinations of three motifs, each 3 to 6 residues long, with the aim of creating short peptides under 20 residues that distill key LLPS-promoting elements from full-length sequences while retaining their phase separation potential.

The “COMBINER” program calculates the extent to which three motifs coexist in DPR sequences by computing the co-occurrence of motif pairs (A with B, and A with C) and representing these as vectors. The sum of these vectors (Vector Score or VS) represents both the number of DPRs in which the three motifs coexist, as well as their symmetry of coexistence. Symmetry in this context refers to the equal occurrence of motif pairs, indicating that motif A co-occurs with motif B as frequently as it does with motif C. This process is repeated for all motif pair combinations, and an average is calculated to produce the Final Score (FS). For a more detailed mathematical explanation of the “COMBINER” algorithm, please refer to the guide in our GitHub repository. This script originated a set of all possible combinations of motif trios, originating small peptides with an average of 12 residues. Additionally, we computed a matrix, using the script “MATRITIAN” that shows the coexistence of the motif trios and their respective FS, that can be found in our GitHub repository.

For the analysis of the designed small peptide sequences, we once again used localCIDER Version 0.1.18, focusing on the parameters Fraction of Charged Residues (FCR), Net Charge per Residue (NCPR), Kappa (κ), Mean Hydropathy and the Fraction Of Disorder Promoting Residues. The program utilized in this step, “CIDER FOR PEPTIDES”, is available in our GitHub repository.

### 3. Experimental validation of designed peptides

#### 3.1 Materials

The peptides were purchased from Genecust (France) with 98.0% purity.

PBS Tablets were obtained from ThermoFisher. Pluronics was purchased from Panreac AppliChem. The microscopy material was obtained from Avantor and Zeiss. The 3’,6’-dihydroxy-6-isothiocyanatospiro[2-benzofuran-3,9’-xanthene]-1-one (FITC) compound was purchased from Sigma-Aldrich.

#### 3.2 Trifluoroacetic acid (TFA) Removal of the Peptides

The peptides were dissolved in a 10mM HCl solution, incubated for 30 minutes, and subsequently lyophilized using a LabConco Freeze-Dryer. This process was repeated three times in total to ensure efficient removal of TFA. The final TFA content was verified to be below 1%.

#### 3.3 Evaluation of LLPS Propensity and Partitioning Experiments

Lyophilized peptide powders were reconstituted in 1mL of distilled water and vortexed until complete dissolution was achieved, resulting in transparent solutions. This process was repeated to obtain three different peptide concentrations: 1 mg/mL, 5 mg/mL, and 10 mg/mL. Droplet formation was induced by mixing 60μL of each peptide solution with 240μL of PBS. ^17^ The sample was incubated for 1 hour at 37°C (±1°C) and left at room temperature (23°C ± 2°C) for 3 hours. To investigate the encapsulation of guest molecules within the condensates, we performed partitioning experiments with the guest molecule FITC (Fluorescein isothiocyanate). Droplet formation was once again induced by mixing 60μL of each peptide solution with 240μL of PBS. The sample was incubated for 5 minutes at 37°C (±1°C), followed by the addition of 1mM FITC. The sample was incubated for 1 hour at 37°C (±1°C) and left at room temperature (23°C ± 2°C) for 3 hours.

#### 3.4 Turbidity Measurements

Turbidity measurements were conducted using a Tecan Infinite M Nano instrument. Absorbance at 600 nm was recorded every 5 minutes for a total duration of 60 minutes. All measurements were carried out at a temperature of 37°C (±1°C). The relative turbidity values reported represent triplicate measurements and were calculated using the formula τ relative % = 100 - T% = 100 - [100% × 10 ^− A600nm^], where A600 is the absorbance at 600nm. A well containing an equivalent volume of buffer solution served as the blank.

#### 3.5 Condensates Imaging

For brightfield images, a Leica DM6000B upright microscope equipped with an Andor iXon 885 EMCCD camera was used. The MetaMorph V5.8 software was employed to control the microscope, and the images were acquired using a 63x 1.4 NA oil immersion phase-contrast objective (Leica). Image analysis was conducted using ImageJ/FIJI 1.54f. ^32^

Confocal images were acquired using a Zeiss LSM 880 point scanning confocal microscope equipped with photomultiplier tube detectors (PMTs) and a gallium arsenide phosphide (GaAsP) detector. To visualize individual condensates, 10μL of the peptide solution was added to Pluronic-functionalized glass slides. Condensates were imaged using a 63x Plan-Apochromat 1.4 NA DIC oil immersion objective (Zeiss) with laser lines at 405nm and 488nm, and appropriate spectral separation for FITC. The Zeiss Zen 2.3 (black edition) software was used to control the microscope, adjust spectral detection for the excitation/emission of the fluorophores used (following manufactures recommendations). Imaging was performed with 1-2% laser intensity for all lasers and a gain between 500 and 650. Image analysis was conducted using ImageJ/FIJI 1.54f. ^32^

#### 3.6 Fluorescence Recovery After Photobleaching (FRAP)

Sample preparation was performed as described in section 3.3. For fluorescence recovery dynamics assessment, a pre-bleached image of the condensates was acquired using 488 nm laser line excitation and emission collected with a GaAsP detector, at 2% power. Subsequently, the target condensates were bleached using the 488 nm laser line at maximum power for 0.45 to 0.93 seconds. The subsequent recovery of the bleached area was recorded using the same acquisition parameters as the pre-bleached image, with a total recovery time of 45 seconds. The final FRAP recovery curves represent the average of three recovery curves collected from n=3 separate droplets. Correction for photobleaching, normalization, and averaging were performed using ImageJ/FIJI. ^32,33^

## Results and Discussion

### 1. Deciphering Amino Acid Motif Patterns in Phase-Separating Proteins

Our first objective was to investigate whether there is a bias of amino acids and peptide motifs in terms of composition, patterning (specific arrangement of residues or motifs), and frequency (repetition of peptide motifs along a sequence) in Phase-Separating Proteins (PhSePs), regardless of their specific function or controlled environments. To this end, we analyzed a cohort of 178 distinct PhSePs, collated from the LLPSDB ^26,27^ and PhaSePro ^28^ databases. These proteins were classified according to their functions into six families: RNA binding, DNA binding, Chromatin binding, Regulation (which includes activators, repressors, and other processing-related proteins), Hydrolases, and Structure (provide support and integrity to biological structures) proteins. The distribution of proteins across these categories was 79, 42, 21, 27, 17, 10, respectively. To ensure statistical validity of our results, we additionally created a negative control database, comprised of 208 proteins with a less than 20% probability of undergoing spontaneous LLPS (referred to as non-PhSePs), as validated by the FuzDrop server. This comparative analysis is key to understand the sequence-specific factors that drive LLPS.

#### 1.1. Amino acid enrichment in PhSePs

The amino acid enrichment analysis was performed on PhSePs, encompassing full sequences, Droplet Promoting Regions (DPRs), and non-Droplet Promoting Regions (NODPRs). This analysis was also performed on PhSePs grouped by families (RNA binding, DNA binding, Chromatin binding, Regulation, Hydrolases, and Structure), as well as in non-PhSePs. The detailed findings can be found in Fig. S1-S6, which show the results for all individual sequences, while Fig. 2 highlights the averaged obtained data.

**Figure 1.**
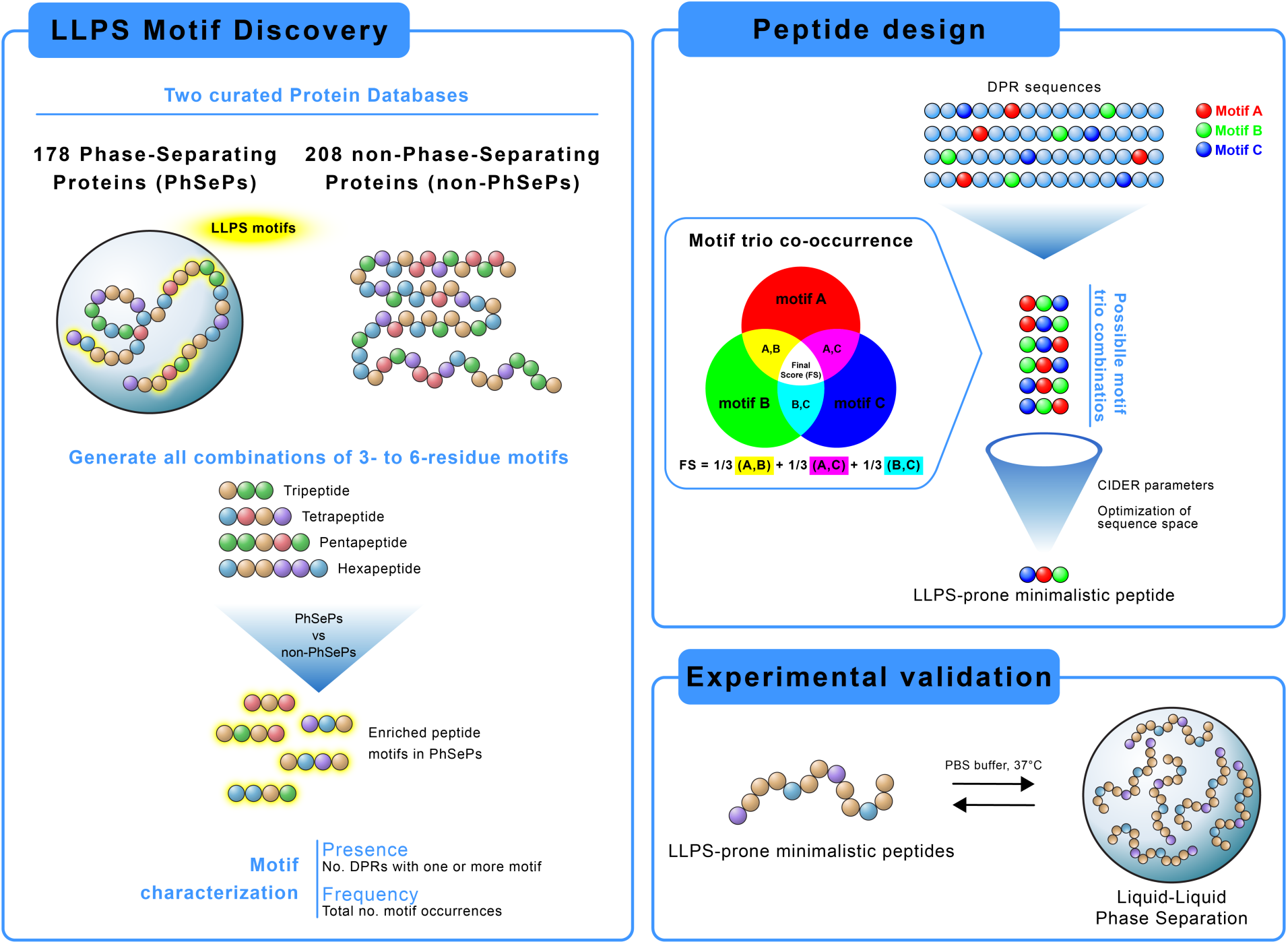
Graphical abstract showcasing our computational approach for LLPS motif discovery and peptide design. The workflow includes: (1) Analysis of Phase-Separating Proteins for identification and characterization of significant peptide motifs (2-6 residues), (2) Data-driven design of minimalistic LLPS-prone peptides by motif co-occurrence studies, and (3) Experimental validation through induced LLPS using synthetic peptides.

**Figure 2.**
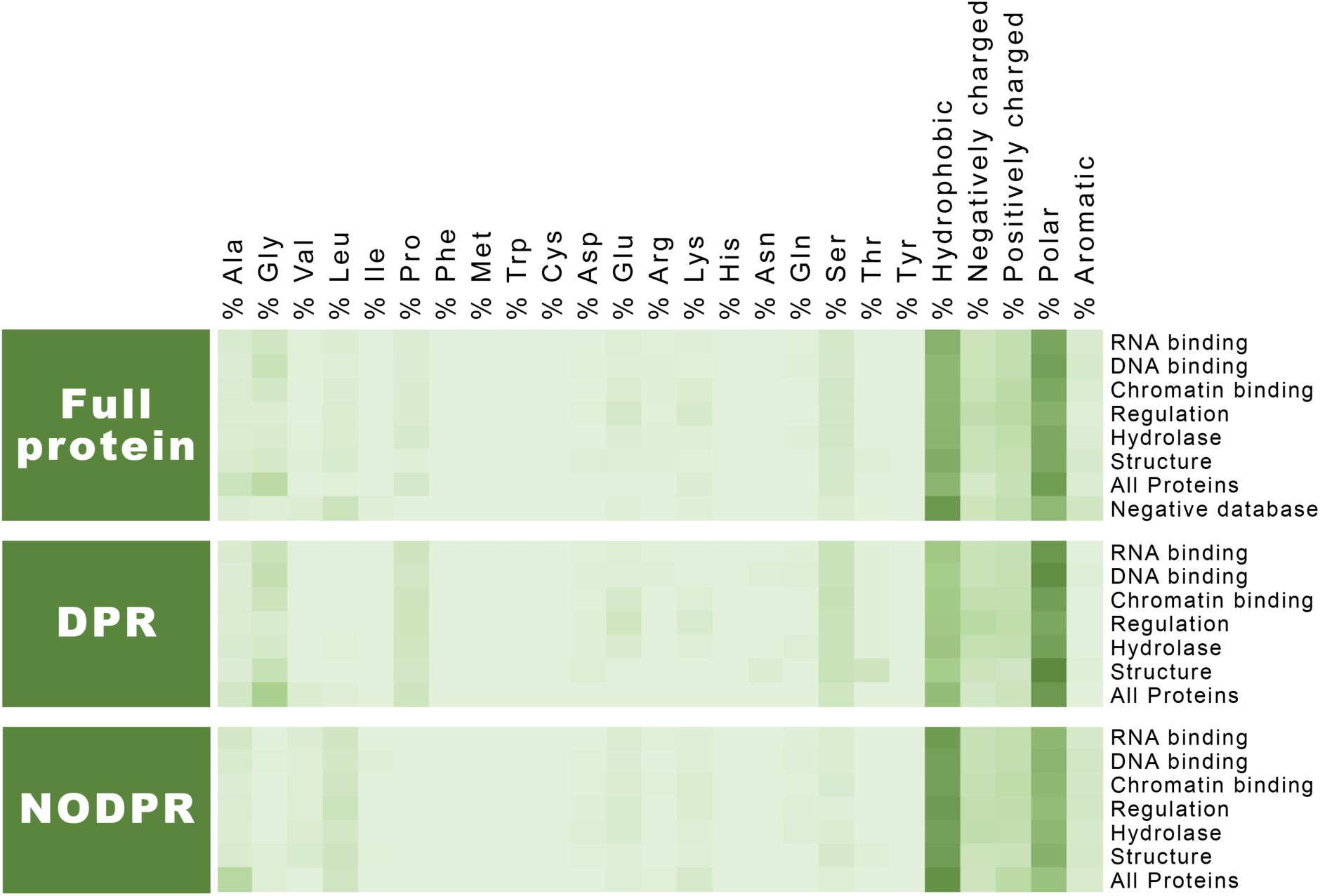
Analysis of amino acids and residue character enrichment in full proteins (including PhSePs database, PhSePs by family, and negative database of non-PhSePs); DPRs (PhSePs database, PhSePs by family) and NODPRs (PhSePs database, PhSePs by family). The color gradient transitions from lighter hues (corresponding to 0%) to darker shades (indicating the maximum value of enrichment in this analysis, corresponding to 50%).

A significant disparity is observed when comparing the distribution of amino acids in the full protein and NODPR regions, against the distributions found within the DPR regions. Notably, the DPR regions show a distinct overrepresentation of Gly, Ser, Pro and Ala. While Gly is not known to directly engage in interactions that lead to LLPS, it is crucial for the dynamics of protein regions in LLPS by maintaining flexible peptide bonds, making it common in disordered regions associated with LLPS. ^4,26^

We also observed a heightened proportion of polar amino acids and a reduced proportion of hydrophobic and aromatic residues within DPR regions. Conversely, NODPRs displayed the opposite trend, characterized by a higher concentration of hydrophobic residues and a lower proportion of polar amino acids. These findings align with prior research indicating that polar residues promote disorder, while small fractions of aromatic and charged residues offer the necessary multivalency for interactions that allow for LLPS. ^3,6^

The unique arrangement of aromatic amino acids in specific patterns is vital for biocondensate formation. They are often interspersed with charged amino acids in mostly polar sequences. While essential for LLPS, excessive aromatic residues can also lead to aggregation. ^5,6^ Their π electron-rich rings facilitate pi-pi stacking, enhancing stability of biocondensates, while charged amino acids promote phase separation through cation-pi interactions. ^2–4,8^

When analyzing the amino acid enrichment in the protein dataset segmented by family, we note an overall similar pattern (Fig. 2, Fig. S3-S6), namely a prevalence of polar amino acids and the reduced proportion of hydrophobic and aromatic residues in DPR regions.

RNA binding proteins have a higher prevalence of polar amino acids, likely to interact with highly charged RNA molecules. Conversely, Chromatin binding proteins are rich in Lys, a positively charged residue that allows for ionic interactions with negatively charged DNA and histones, crucial for chromatin-protein complex formation and regulation of chromatin compaction. ^34^

Structure proteins are enriched in Gly, Ser, Pro, Ala and Val. Gly provides flexibility and stability, properties common to proteins such as collagen, elastin and keratin. ^35–39^ Val contributes to integrity and hydrophobic interactions within the protein, and is found in high proportions in elastin, titin and fibroin, ^36,40,41^ while Pro in enriched in collagen and elastin, ^35,36,38,39^ and Ser in keratin and fibroin. ^37,41^ Moreover, Val and Ala are components of repeating motifs found in elastin and fibroin. ^38–41^

The enrichment of Ala, Asp, and Thr in Hydrolases is logical, since Asp and Thr are crucial catalytic residues while Ala aids in maintaining enzyme flexibility and accessibility, essential for conformational changes and substrate interactions in catalysis. ^42^ The prevalence of these residues in enzymes undergoing LLPS suggests a potential link between amino acids involved in LLPS and catalysis, which is plausible to hypothesize given that LLPS is known to enhance catalytic activity in natural systems. ^43,44^

The analysis of the negative database reveals completely distinct trends, including a higher average of hydrophobic and aromatic residues and a lower frequency of polar amino acids. Table 1 underscores that the key residues enriched in PhSePs, such as Gly and Pro, are absent in non-PhSePs, with the exception of Ser. Additionally, non-PhSePs show an overrepresentation of Leu residues, a pattern not observed in PhSePs.

**Table 1.**
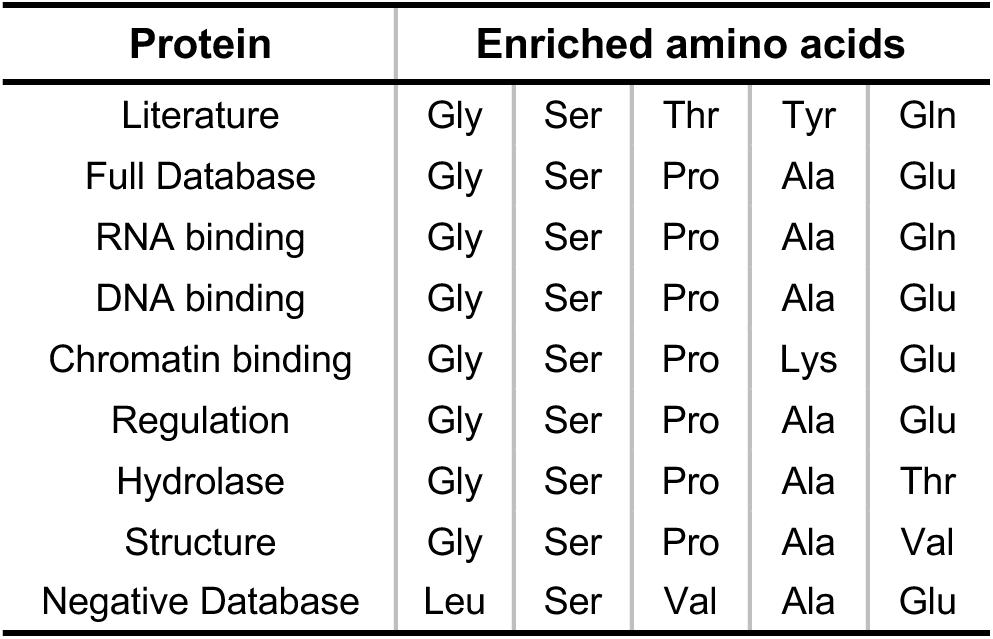
Top enriched amino acids influencing LLPS. Represented is literature-based commonly reported amino acids ^1–4,29^, our database analysis of the DPR regions of 178 PhSePs, protein family-based analysis of DPR, and negative database analysis on non-PhSePs.

Having conducted a comprehensive analysis of the amino acid characteristics in DPRs and NODPRs, we sought to explore whether we could differentiate between the properties of the primary sequences of DPRs and NODPRs. To accomplish this, we utilized the CIDER (Classification of Intrinsically Disordered Ensemble Regions) server, a powerful tool developed by the Pappu lab. ^23^ CIDER is designed for the analysis of parameters associated with the primary sequence of IDPs, thus giving insights about the behavior of unstructured ensembles. ^23,29–31^ For the analysis of DPRs and NODPRs, we focused on five key metrics used in the comprehensive work of Ginell and Holehouse, ^31^ who calculated these across all IDRs in the human proteome. The parameters are the Fraction of Charged Residues or FCR (proportion of charged amino acids), the Net Charge per Residue or NCPR (overall charge of the sequence, considering positive and negative charges), Kappa or κ (quantifies the extent of charge mixing along the sequence), Mean Hydropathy (reflects the hydrophobicity of a sequence) and the Fraction of Disorder-Promoting Residues (proportion of amino acids predicted to be disorder-prompting, here encompassing Ala, Arg, Gly, Gln, Ser, Pro, Glu, and Lys). ^23,30,31^ The resulting graphs plots obtained from analyzing the DPR and NODPR sequences using these parameters are presented in Fig. S7.

Both charge-based parameters (FCR and NCPR) show similar trends between DPRs and NODPRs, falling within the range observed for human IDRs. ^31^ This aligns with our previous amino acid analysis, which revealed no significant difference in the overall charge content among full PhSePs, DPRs, and NOPPRs. Although DPRs may contain repeating and enriched motifs with charged amino acids, we propose that NODPRs exhibit a similar charge content without the same repetitive patterns, resulting in comparable overall charge percentages. Consequently, these charge-based parameters alone are insufficient for distinguishing between DPRs and NODPRs.

Charge clustering and segregation are recognized as important factors in LLPS, with clustered charges promoting such phenomenon. ^23,26,45^ Therefore, it is not surprising that the κ parameter, which measures charge distribution, differs slightly between DPRs and NODPRs. For DPRs, the κ value falls within the range of human IDRs, ^31^ while for NODPRs, it does not. The higher value for DPRs (where κ closer to 0 indicates more distributed charge and closer to 1 indicates more clustered charge), indicates a more clustered presence of charged residues. This parameter provides a means to distinguish DPRs from NODPRs and further associate DPRs with known characteristics of sequences prone to LLPS.

Hydropathy is a measure of the relative hydrophobicity or hydrophilicity of the amino acids in a protein sequence. ^46^ This metric can be used to differentiate DPRs from NODPRs, where hydropathy values for DPRs falls within the range observed for human IDRs (corresponding to lower hydrophobicity), while the value for NODPRs exceeds the upper limit of this range. This is corroborated by the literature, that highlights that IDPs are known to have lower hydrophobic content compared to structured proteins. ^23,31,47^

DPRs also exhibit a higher proportion of residues involved in LLPS, with a value within the range observed for human IDRs. ^31^ In contrast, NODPRs have a lower proportion of such residues, with a value outside the IDP range. This clear distinction in LLPS-prone residue content provides a robust metric for differentiating between DPRs and NODPRs.

Our analysis aligns with the known literature regarding amino acid enrichment patterns in PhSePs. Although minor variations exist among different protein families, such differences are consistent with their specific functionalities (e.g., catalytic residues in Hydrolases), indicating that the propensity of a protein to undergo LLPS may also depend on its specific family context. Regarding sequence differences between DPRs and NODPRs, several metrics usually used for IDPs can be used to distinguish between these. The main differentiating factors include those related to charge patterning, hydrophobic content, and proportion of residues associated with disorder.

#### 1.2. Motif Discovery in PhSePs

Amino acid patterning, which refers to the arrangement or organization of specific amino acids in motifs, is also fundamental in LLPS. ^1–3,5,7,8,29,48^ After the analysis of DPR and NODPR regions of all PhSePs, it became evident that NODPR regions exhibit a prominently randomized distribution of residues. In contrast, DPR regions tend to display significantly more consistent and enriched patterns.

To investigate which patterns are enriched in our DPR database, we comprehensively examined all peptide motifs combinations ranging from three to six amino acids in length, by computing their presence (number of distinct DPRs with at least one instance of a motif) and frequency (total occurrences across all DPRs, including repeats within sequences), represented in Fig. 3A. To ensure the statistical validity of the obtained values, we repeated the same process using our negative database of non-PhSePs. The addition of a negative database for data correlation is key to understanding the sequence-specific factors that drive LLPS, as we aim to study motifs enriched in proteins that undergo phase separation, rather than in proteins in general.

**Figure 3.**
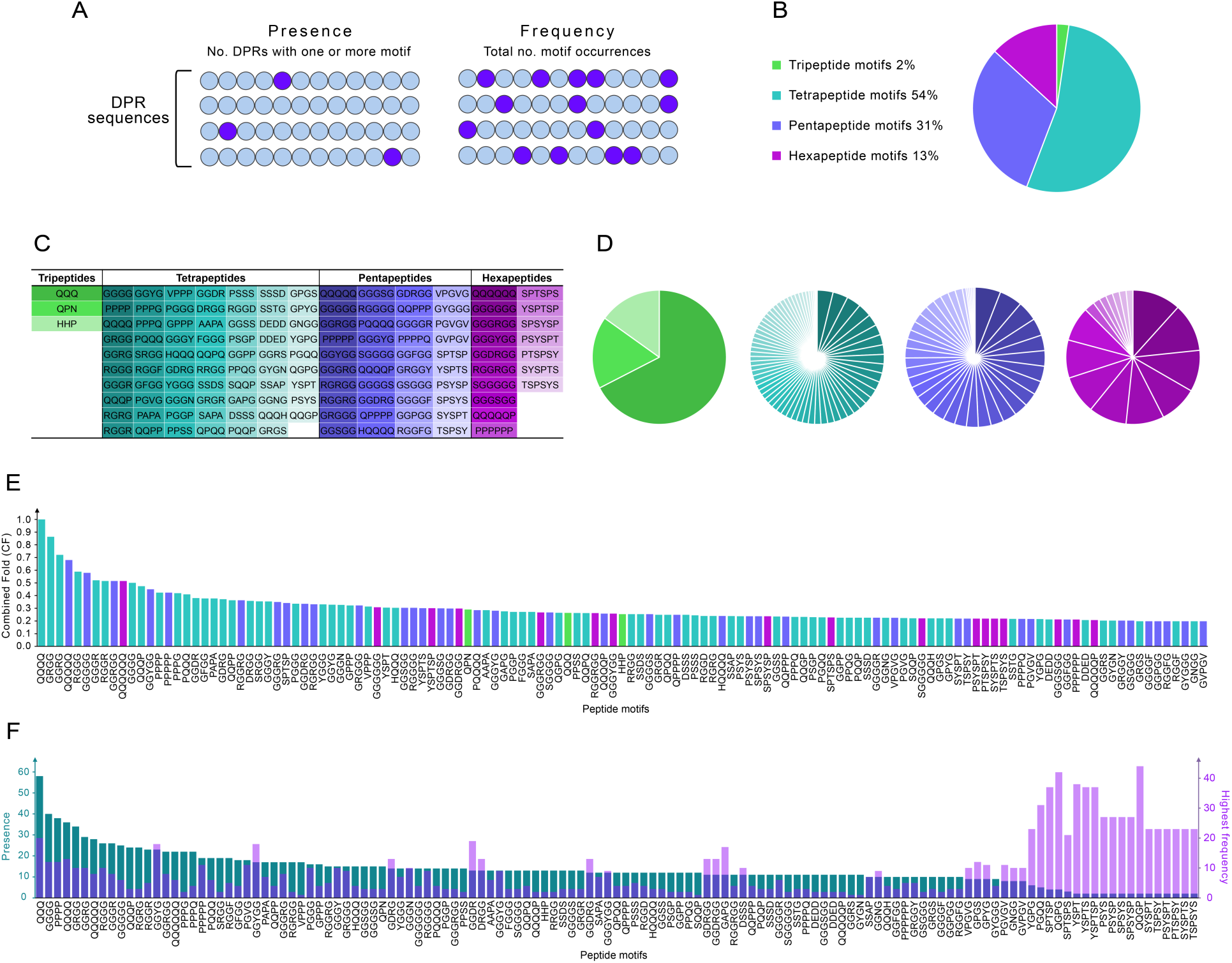
Motif discovery analysis of PhSePs. (A) Schematic representation of the Presence and Frequency parameters, (B) Distribution of the 129 discovered peptide motifs by length, ranging from tripeptides to hexapeptides, (C) Listing of the 129 identified peptide motifs, (D) Proportion of enriched peptide motifs based on their presence values depending on peptide lengths, (E) Distribution of peptides based on their CF values, (F) Presence values of the motifs organized in descending order, along with their highest frequency value observed in any DPR.

Motif significance was assessed using a combined fold score (CF) that equally weighted the normalized presence fold (PF = Presence in DPRs of PhSePs/Presence in non-PhSePs) and normalized frequency fold (FF = Frequency in DPRs of PhSePs/frequency in non-PhSePs) of each motif. Both PF and FF, as well as CF have normalized values between 0 and 1. Motifs with a CF ≥ 0.2 were considered, resulting in 129 motifs for further analysis, as illustrated in Fig. 3C.

From the 129 motifs identified, 36% were found to be embedded within others (e.g., QQQQ in QQQQQ, GRGG in GGRGG), with the majority of these being embedded in only 1-2 other motifs. We chose not to eliminate any of these instances because we consider that even a single amino acid addition may significantly impact the LLPS behavior we are studying.

To validate that the enrichment of these 129 motifs in DPRs differs from that in NODPRs, we also analyzed motif presence and frequency in the NODPRs sequences. Our motifs appear, on average, in 1 out of 702 NODPRs (with a mean frequency of 1), while they are found in 14 out of 712 DPRs (with a mean frequency of 31), as shown in Table S1.

#### 1.3 Characterizing Motif Composition and Diversity in PhSePs

Among the resulting 129 peptide motifs, the length distribution includes 2% tripeptides, 54% tetrapeptides, 31% pentapeptides and 13% hexapeptides (Fig. 3B). This distribution suggests that tetrapeptides might serve as the fundamental unit for LLPS and could represent an optimal balance between length and interaction potential, contributing to the stability of phase-separated states. ^14,49^ In Fig. 3C and D, the distribution of the 129 peptide motifs is illustrated regarding their presence in the DPR sequences and organized by peptide length. Based on average PF and FF values, tetrapeptides are the most present, followed by tri- and pentapeptides. While hexapeptides are less common overall, they appear with higher frequencies in DPR sequences, followed by tetra- and pentapeptides. This indicates that shorter peptides are generally more enriched in the DPR database, while longer peptides, though less abundant, appear to show more repetitions in fewer sequences. Fig. 3E further shows the distribution of discovered motifs based on CF values.

Among the discovered enriched motifs, individual amino acid trends are observed. Gly is notably overrepresented, appearing in 59% of motifs, followed by Pro (47%), Ser (29%), Arg (23%), Gln (20%) and Tyr (19%). When comparing the amino acid enrichment in the peptide motifs with the overall enrichment in DPR sequences in Fig. 1, we observe that Gly, Pro, and Ser are the top three residues in both analyses, while Arg, Gln, and Tyr showed no significant enrichment in DPR sequences. In fact, Tyr ranks among the bottom five amino acids present in DPRs. This suggests that specific arrangements of amino acids within short peptide motifs, composed of polar (Gly, Ser, Gln) and hydrophobic (Pro) residues and interspersed with charged (Arg) and aromatic (Tyr) residues, may be the key for a minimalistic approach to LLPS of small biomolecules.

Further trends can be found in the set of 129 motifs. Firstly, distinct clusters of homopeptides are present, with notable repetitions of Gln (3 to 6 residues), Pro (4 to 6 residues), and Gly (5 to 6 residues), which are characteristic of low-complexity regions (LCRs) within IDPs. Gln and Gly are significantly represented within LCRs, ^6,7^ while Pro is believed to contribute to the nuanced structural dynamics of IDPs. ^50,51^ The enrichment of these homopeptides in PhSePs compared to non-PhSePs suggests their key role in enhancing the flexibility and dynamics critical for phase separation, highlighting their greater importance in proteins that undergo LLPS. Interestingly, our analysis revealed not only such clusters but also similar sequences with additional residues interspersed or flanking these clusters, such as QQQP, SGGGG, HQQQ, and GGGR. These additional residues possibly fine-tune the LLPS propensity of proteins by providing specificity, modifying interaction strengths, or influencing the physicochemical properties of the resulting biomolecular condensates.

Charged amino acids are known for facilitating a variety of interactions both within and between proteins. ^7,48^ In our analysis, several motifs contain charged residues, primarily the positively charged Arg and negatively charged Asp. Interestingly, Arg consistently appears with Gly, typically in a 1:2 to 1:5 ratio (e.g., RGGF, GGRS, GRGGY, GGGRGG). A subset of Arg containing motifs also include the oppositely charged Asp (e.g., DRGG, GDRG, GGDRGG). Motifs containing His are always paired with either Gln or Pro (e.g., HHP, HQQQ). Remarkably, entirely negatively charged motifs (with Asp and Glu) can be found in our set (e.g., DDED, DEDD). Motifs with Asp often co-occur with Gly or Ser (e.g., GGDR, SSDS). Such patterns of charged amino acids with specific residues appears to be important for PhSePs. These recurring combinations likely play key roles in facilitating electrostatic interactions and hydrogen bonding at protein interfaces. The prevalence of Gly in many of these motifs may provide conformational flexibility to optimize charged residue positioning for intermolecular contacts.

The RG/RGG motif, which is a GAR (Glycine- and arginine-rich) motif, is arguably one of the most known motif pattern related with LLPS. ^11,19,52–54^ This sequence, prevalent in RNA-binding proteins (RBPs), is involved in protein-RNA interactions through Arg-Gly mediated cation–π, π– π, hydrogen-bonding, and electrostatic interactions. ^11,53^ By driving LLPS of several proteins in cells (i.e. FUS, EWS, TAF15, etc.), the RG/RGG motif allows for the formation of membraneless organelles that aid in the regulation several cellular functions, including ribosome biogenesis and mRNA regulation. ^47,48,50^ Interestingly, the RGG motif itself did not appear among the motifs we discovered. This is due to it being equally present in the non-PhSePs database, leading to a lower fold value compared to motifs that are more enriched in PhSePs. However, we did identify several variations of patterns where the RGG motif was incorporated (a total of 19 motifs), such as DRGG, GRGG, RGGF, GRGGY, GGDRGG, and even a double variation of the motif, RGGRGG. This finding suggests that, while RGG domains can undergo LLPS, the context of specific additional amino acids - consistently identified in our results as Asp, Arg, Gly, Ser, Phe, Tyr, or combinations of up to three of these - may significantly influence condensate formation in synthetic systems.

Aromatic residues, particularly Tyr and Phe, play crucial roles in protein-protein interactions in LLPS, despite their relatively low abundance previously reported. Such residues engage in π-π stacking and cation-π interactions, contributing to multivalent interactions essential for the formation of biomolecular condensates. ^4,5,8,55,56^ In our discovered motifs, the prevalence of Gly residues surrounding the aromatic amino acids is striking, with Tyr-containing motifs (e.g., GGGY, GRGGY, GYGN) and Phe-containing motifs (e.g., FGGG, GGFGG, GGGGF, RGGFG) being particularly abundant. The number, composition, and patterning of aromatic residues are extremely important in condensate formation and aggregation inhibition, as well as in determining the properties of the droplet, including diffusion rates, biochemical stability, and material phase. ^5,56^

YGG motifs can form dense droplets in the context of RNA- and DNA-binding proteins such as hnRNPA2, ^19^ FUS and RBM3. ^52^ Such motifs facilitate protein-protein and protein-nucleic acid interactions, with Tyr engaging in π-π stacking and cation-π interactions, while Gly provide flexibility. Similarly, to the RGG motif, we did not find the isolated form of the YGG motif, but several variations that incorporated additional Gly residues, including YGGG, GGYG, GGYGG, GYGGG, and GGGYGG. Furthermore, our analysis uncovered several other motif variations containing the aromatic Tyr, specifically GPYG, YGPG, and GYGN. Of particular interest, we identified the GRGGY motif that appears to be a hybrid of the RGG and YGG patterns. Again, we report that although the YGG motif unit it known to undergo LLPS, more complex variations with additional residues and patterns might further elucidate on the LLPS mechanism and dynamics.

We identified the VPGVG short peptide, as well as adjacent motifs, VPPP, PGVG, PGVGV, and GVPGV, which show partial matches with motifs commonly found in disordered elastins, proteins known for their phase transition capabilities. ^38–40^ These proteins typically contain repetitive motifs that allow for their phase separation behavior, such as the canonical pentapeptide VPGVG, as well as variations like VPGG and GVGVP. ^36,38–40^ In these motifs, Val residues provide hydrophobicity, Gly offer flexibility, and Pro introduces conformational constraints. ^36,39,40^

One notable category of small peptides in our database consists of 7 motifs (YSPT, SPTSP, TSPSY, YSPTS, PTSPSY, SPTSPS and YSPTSP) and 6 motifs (DRGG, GDRG, GGDR, GDRGG, GGDRG and GGDRGG). These peptide motifs are embedded within YSPTSPSY and YGGDRGG, which are known periodic repeats present and enriched in PhSePs (RNA polymerase II and TAF15, a member of the FET family of RBPs) that form condensates. ^57–59^

Repetitive motifs in PhSePs are essential for LLPS, as their recurring patterns within a protein sequence significantly modulate phase separation. It is thought that a uniform distribution of specific residues induces LLPS, whereas the absence of this particular distribution can lead to the formation of amorphous precipitates. ^2,5,8^ Hence, we aimed to examine the presence of motifs (number of DPRs containing a motif), motif frequency (occurrences across all DPRs, including repeats within sequences), as well as compare the interplay between the two parameters in the discovered motifs. The resulting data in Fig. 3F displays the overall presence of the motifs organized in descending order, alongside their highest frequency value observed in a single DPR.

The analysis reveals an equilibrium between the presence and frequency of motifs in PhSePs. On average, motif presence is 3 to 4 times higher than frequency, suggesting a balanced distribution of these elements across different protein, with some exceptions visible, as is the case for the motifs GGYG, GGYGG, GDRG, DRGG, GGDRG, GGGYGG, GDRGG, GGDRGG, GAPG, DSSS, SSAP, GGNG, and GVPGV. This suggests that these motifs, when present in a protein, tend to occur with higher repetitions within the same sequence.

Conversely, the right side of the graph (Fig. 3F) highlights motifs that, despite their limited presence across proteins, display high repetitiveness within specific sequences. Notably, partial sequences of the periodic repeat YSPTSPSY demonstrate some of the highest frequency values among the motif list (appearing between 23 and 38 times within a single DPR sequence). Additional highly repetitive motifs YGPG, PGQQ, QGPG, and QQGP, have frequencies ranging from 23 to 44 repetitions in a DPR. Higher motif repetitions in these proteins might imply a distinct mechanism in LLPS, where multiple copies of specific motifs are more crucial than in typical PhSePs. ^60^

Although most of the enriched peptide motifs align with previous results, thus validating our methods, we have also identified several new motifs. These include HHP, QPN, PAPA, AAPA, SAPA, GAPG, GPGS, QQPP, QGPG, PSGP, PPQG, PPSS, SSDS, SSAP, DSSS, DDED, and DEDD. These newly identified motifs could potentially play a significant role in phase separation, thereby expanding our understanding of the sequence determinants of this phenomenon.

#### 1.4 Family-specific motif variations

RNA-binding proteins (RBPs) are the most extensively studied proteins undergoing LLPS. ^11,19,52–54^ Research has revealed that specific sequence motifs drive LLPS by mediating multivalent interactions, aligning with the stickers and spacers model. However, this model has been primarily validated in Prion-like RBPs. ^1,2,5^ To determine if motif enrichment is a general LLPS feature or function-specific, we expanded our analysis to diverse protein families beyond RBPs. We have compared motif enrichments across these families to determine if LLPS-associated motifs are function-dependent or represent a broader, function-independent phenomenon.

We performed a similar analysis as done previously, now on the protein family sub-databases, encompassing peptide motif discovery and characterization, while using non-PhSePs as a negative control. Fig. 4A depicts motif length distribution, while Fig. 4B illustrates motif presence in descending order, alongside their highest frequency within a DPR.

**Figure 4.**
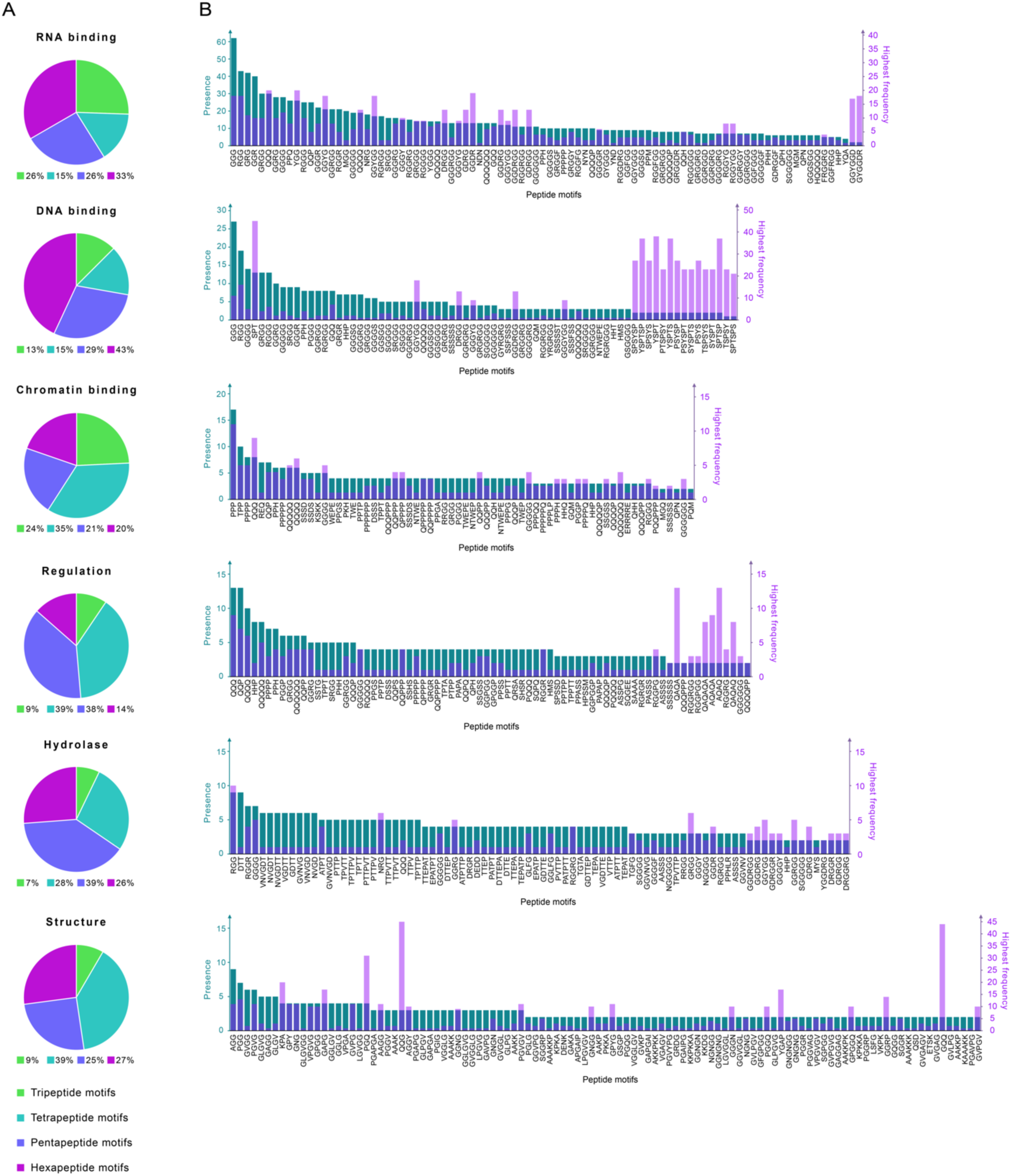
Motif discovery analysis of PhSePs segregated by family. (A) Distribution of the discovered peptide motifs by length, ranging from tripeptides to hexapeptides, (B) Presence values of the motifs organized in descending order, along with their highest frequency value observed in any DPR.

Comparative analysis of motif length distribution across protein families revealed distinct patterns, with tripeptides showing significant enrichment (7-26%) compared to only 2% in full PhSePs, while tetrapeptides, dominant in PhSePs (54%), were underrepresented in families (15-39%). Pentapeptides maintained relatively prevalence, while hexapeptides exhibited notable enrichment compared to PhSePs (13-72%).

Regarding distinct families, RNA and DNA binding proteins favor longer penta- and hexapeptides, Chromatin binding proteins show slight preference for shorter tri-/tetrapeptides, while Regulation proteins prefer mid-length tetra- and pentapeptides. Hydrolases are enriched in pentapeptides, followed by tetra- and hexapeptides, and Structure proteins favor tetrapeptides, closely followed by penta- and hexapeptides.

Analysis of amino acid composition in discovered motifs across protein families revealed both conserved trends and function-specific patterns. While the general amino acid enrichment trend persisted, specific amino acids and their ratios vary depending on the functional context. The ubiquity of Gly and Pro suggests their universal role in phase separation, independent of protein function; in contrast, Tyr and Ser, which are enriched in general PhSePs motifs, showed limited presence in family-specific motifs. The nuanced differences in other enriched amino acids likely reflect context-dependent molecular interactions crucial for each family’s specific function.

Three motifs (HHP, GRGG, and GGGGG) are present in five out of six families, excluding the Structure family, while several others (e.g., QQQ, PPH, GGGG, GGRG) appear in four families. The prevalence of Gly-rich and Pro-containing sequences highlights the role of structural flexibility in LLPS, indicating a common mechanism across cellular functions. Notably, 40-50% of motifs are unique to binding and Regulation families, 73% to Hydrolases, and 99% to the Structure family, suggesting that some motifs contribute to LLPS in general, while others seem to be intrinsically linked to specific functions. The unique motifs of each protein family are listed in Table S2.

In RNA binding PhSePs, a significant number of motifs are RG/RGG-adjacent patterns, ^11,19,52–54^ including RGGGG, FRGGRG, GGRGGD, GRGGDR and GRGGY, as well as YGG and FGG patterns, ^19,52^ such as YGG, RGGYGG, GGYGGD and GGFGG. We also identified clusters of Gln and Gly, along with versions with additional amino acids like HQQQQQ, GGGGR, and GGGGGF. The discovered motifs show an overall balance between the presence and frequency parameters (Fig. 4B), with some outliers (e.g., QQQ, YGG, GGYG, GGYGG) exhibiting higher frequencies than presence. Interestingly, motifs such as GGYGGD and GYGGDR have low presence but extremely high frequency, appearing 17 and 18 times in a sequence, respectively.

In the DNA binding category, we observed YSPTSPSY-derived motifs (YSPTSP, SPSYS, PSYSP), ^57,58^ as well as RG/RGG patterns (RGRGGG, SRGGG, YRGRGG). Clusters of Gly, Ser, and Gln acid residues were also prominent in our analysis and, notably, we observed many motifs composed solely of Gly and Ser (i.e. GGSGG, GSGGG). These Ser/Gly-rich patterns may be related to the [G/S]Y[G/S] domain found in the LCRs of the FUS protein.^19,61,62^ Analysis of Fig. 4B reveals that motif presence and frequency are balanced, similar to the RNA binding proteins, although with slightly lower frequency values. A notable exception is the motifs related to the periodic repeat YSPTSPSY, which appear to be highly repetitive, with frequencies 11 to 23 times higher than presence values.

Several motifs with positively charged residues were found in Chromatin binding proteins, such as HQQ, PKH, PPPH, KSKK and ERRRRE. Chromatin has been shown to undergo LLPS, driven by the interactions with positively charged histone tails, a chromatin binding protein. ^63,64^ We also found many motifs enriched in Gln and Pro, likely due to their role in promoting multivalent interactions important for chromatin-associated protein LLPS. ^64,65^ However, the specific contribution of Gln/Pro-rich motifs to chromatin binding in LLPS contexts is not well-established in current literature. The motifs found in Chromatin binding proteins display a close ratio between presence and frequency parameters.

The motifs enriched in the Regulation family, are Gln-rich sequences (PQQQ, RQQQQ, QQPQ, QAQAQ), are associated with transcriptional activation domains, ^66–68^ Gly-rich motifs (GGPGG, GPGGP, RGGPG, RGRGR) are observed in dynamic proteins assemblies involved in transcriptional regulation and signal transduction. ^18,69^ Pro-rich motifs (PAPAP, PPTPP, PPTT, PPSS) mediate crucial protein-protein interactions, ^20,66,68,70^ and Ser-rich sequences (SPSSD, SSTG, SHSR, ASSPG) are phosphorylation sites for activity regulation. ^68,71,72^ The repetitive nature of amino acids in motifs (QAQAQA, AQAQA, PAPAP) may suggest a contribution to the structural flexibility required for dynamic regulatory interactions. ^68,73^ Interestingly, most motifs in Fig. 4B are generally more present than frequent, except for those on the right side of the graph (e.g., QAQA, AQAQ), which consist solely of Gln and Ala and are 4 to 6 times more frequent than they are present.

Enriched motifs in Hydrolases contain amino acids typically found in catalytic triads of hydrolases, such as the His in PPHLR, or Asp and Glu in DEDD, DTTEP, VGDTTE, and GDTTEP. ^42,74^ Aromatic residues, in motifs like GGLFG and TGFG, MYS, can be significant for protein-protein interactions, which are important in both enzymatic function and LLPS behavior. ^75,76^ We observed a high proportion of motifs enriched in Pro, Val, and Thr that seem to have a repetitive nature (PTTPVT, TPTTPV, TTPVTT, PTTPV, TTPVT, PVTTP, among others), indicating they may be part of larger elements that contribute to the protein’s architecture and phase separation. ^3,29^ By analyzing Fig. 4B, we note that motifs in the Hydrolase family are 3 to 6 times more present than frequent. However, there are outliers with higher frequencies, such as GRGG, GGDRGG, and GGYGG.

In the Structure family, we observe a prevalence of Val/Pro-enriched motifs, including GAVPG, GVLPG, GVPG, VPGA, and VPGVG, among others. These sequences align closely with the motifs VPGVG, VPGG, and GVGVP, characteristic of elastin-like polypeptides (ELPs). The abundance of these motifs highlights their essential role in providing elasticity, overall structure, and ability to undergo phase transitions. ^36,38–40^ In Structure proteins, motifs are both highly present and frequent, with QQG and GQQ motifs showing exceptionally high frequencies of 45 and 44 occurrences, respectively.

Our analysis revealed diverse LLPS-associated motifs across protein families, indicating unique preferences likely shaped by specific functional requirements, thus underscoring the complex relationship between sequence, function, and phase separation behavior. The varied distribution of motif lengths, presence and frequency values across families suggests distinct mechanisms for LLPS.

### 2. Minimalistic peptide design based on motif co-occurrence

Our design of minimalistic peptide sequences with LLPS propensity relied on the co-occurrence of enriched motifs, which can create synergistic effects that potentially modulate a biomolecule’s ability to undergo LLPS.

We explored all possible combinations of three motifs from the 129 identified in PhSePs (Fig. 5). For each trio of motifs, we studied their co-occurrence patterns within DPR sequences, by examining how often pairs of motifs appear together (A with B, A with C, and B with C) and the symmetry of their presence (for a more detailed mathematical explanation of our approach, refer to the guide in our GitHub repository). This analysis led to a scoring system (Final score or FS) that reflects both the frequency and balance of motif co-occurrence. A perfect score of FS equal to 100 would indicate that all three motifs are present in all DPRs in equal proportions. Scores closer to 50 might suggest either an asymmetrical presence of a motif pair over another or a limited overall presence across DPR sequences. Scores approaching 0 indicate minimal to no co-occurrence of the three motifs. A matrix was computed showcasing the co-occurrence patterns and respective scores for all possible motif trios, and can be found in our GitHub repository.

**Figure 5.**
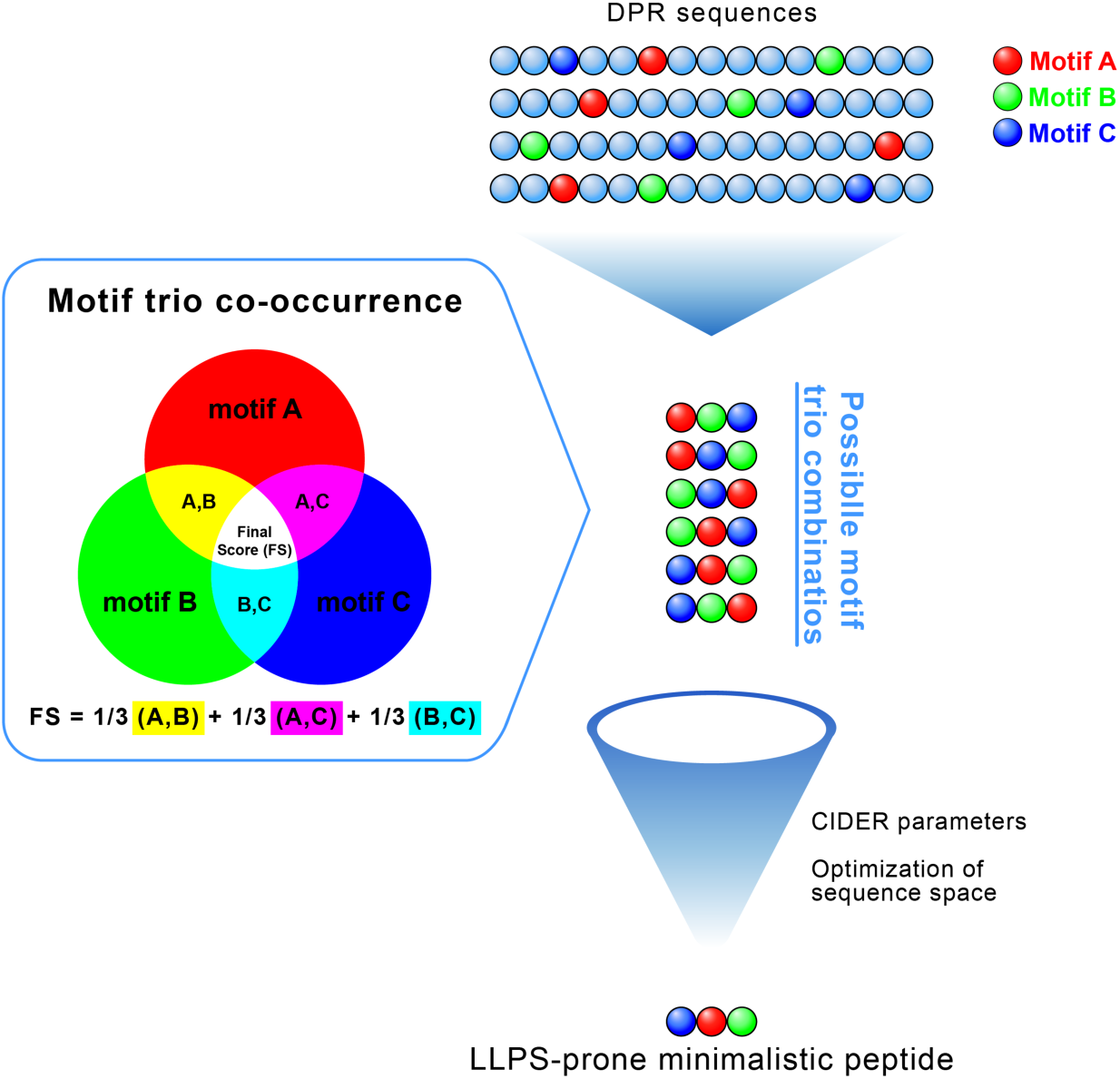
Conceptual approach to peptide design. We analyzed motif trios from the 129 discovered motifs in the context of DPRs, focusing on their presence and symmetry of co-occurrence patterns. All permutations of these motif trios were computed, assigned scores based on their co-occurrence analysis, and ranked by their final scores before further refining them using CIDER parameters and sequence space optimization. This process yielded a list of short peptide sequences with high potential for LLPS.

We subsequently generated all possible permutations of each motif trio, thus yielding a diverse set of short peptide sequences. These initial designs were refined by identifying and merging overlapping regions between adjacent motifs, eliminating redundancies and optimizing the sequence space, resulting in peptide sequences with an average length of 12 residues.

Our analysis of the top 10 short peptides ranked by FS reveals a significant overrepresentation of motifs composed of Gly and Arg residues, followed by motifs rich in Gly interspersed with Phe, Ser, Tyr, Asp, His and Glu. While these amino acids are known to contribute to LLPS through various mechanisms, our unbiased methodology provides novel insights into their precise combinations and patterns.

While the top-ranking peptides were predominantly composed of Gly and Arg, an examination of the top 100 motif combinations reveals a broader spectrum of amino acid compositions and patterns. These include Pro- and Gln-rich sequences (PPPG, PPPPQ, QQPPP, QQQH, QQQQ), as well as Ser-rich (GGRS, GGGGS, GSGGG). This expanded set of motifs underscores the diversity of sequence patterns that may contribute to phase separation properties, suggesting that a wider range of amino acid combinations play significant roles in LLPS beyond the most prominent Gly-Arg pairings.

To refine our selection of designed peptides from the diverse motif trio permutations, we utilized the CIDER server to compare their characteristics with those of known human IDRs, thereby enhancing our understanding of how these motif arrangements align with typical IDP features. ^23,29–31^ We focused on the same key parameters used previously (section 1), drawn from the comprehensive work of Ginell and Holehouse, ^31^ calculated in IDRs of the human proteome. Our list of peptides was thus refined and filtered using the range of values for the parameters FCR, NCPR, κ, Mean Hydropathy and Fraction of Disorder Promoting Residues, and organized according to the FS scores. The distribution of these parameters in the designed peptide sequences is shown in Fig. S8.

We generated a peptide library (Table 2) featuring diverse motifs. To ensure a comprehensive representation of our motif discovery results, we intentionally selected peptides that did not share the same motifs, allowing us to explore a broad spectrum of structural and functional possibilities.

**Table 2.** Peptide sequences obtained by computational approach.

| Peptide | Sequence | Final Score (FS) |
| --- | --- | --- |
| PJ1 | FGGGRGGFGGDRGG | 74.00 |
| PJ2 | GRGGYGGDRGGYGG | 61.26 |
| PJ3 | GYGGGFGGDRG | 60.51 |
| PJ4 | GYGNDRGGSGGGG | 55.26 |
| PJ5 | GPYGDRGGFG | 29.36 |
| PJ6 | PGVGGYGDRGG | 20.90 |
| PJ7 | RGGFVPPPRGGD | 17.87 |
| PJ8 | YSPTSPGGDRGGFG | 14.87 |

We designed eight peptides (PJ1-PJ8) to systematically explore the impact of amino acid composition on phase separation propensity. PJ1, the peptide with the highest FS score from our co-occurrence analysis, serves as a baseline with Gly, Arg, and Phe. Subsequent peptides introduce variations: PJ2 substitutes Tyr for Phe, allowing us to compare the effects of different aromatic residues. PJ3 incorporates both Tyr and Phe, providing insight into the interplay of multiple aromatic amino acids. PJ4 introduces polar residues Asn and Ser, potentially altering the hydrophilicity and hydrogen bonding capacity of the peptide. PJ5 and PJ6 feature Pro and Val, respectively, allowing us to examine the impact of a cyclic amino acid and a hydrophobic residue on disorder propensity. PJ7 stands out with its Pro triplet (Pro-Pro-Pro) and the strategic placement of charged residues (Arg and Asp) separated by Gly, potentially influencing the peptide’s conformational flexibility and stability. Lastly, PJ8 incorporates a partial sequence of the established LLPS-associated YSPTSPSY motif, serving as a benchmark for phase separation propensity. Additional notable differences include varying peptide lengths, from 10 to 14 residues, and different patterns of Gly distribution, which may affect overall peptide flexibility.

### 3. Experimental validation of designed peptides

Using the custom designed peptides, our aim was to evaluate their capability to form condensates under physiological conditions (PBS buffer, 37°C), focusing solely on the inherent ability of the primary sequences to undergo LLPS. To investigate this, we conducted a screening approach using the PJ peptides at three different concentrations (1, 5, and 10 mg/mL). We characterized the resulting samples through optical brightfield microscopy and relative turbidity measurements (Fig. 6).

**Figure 6.**
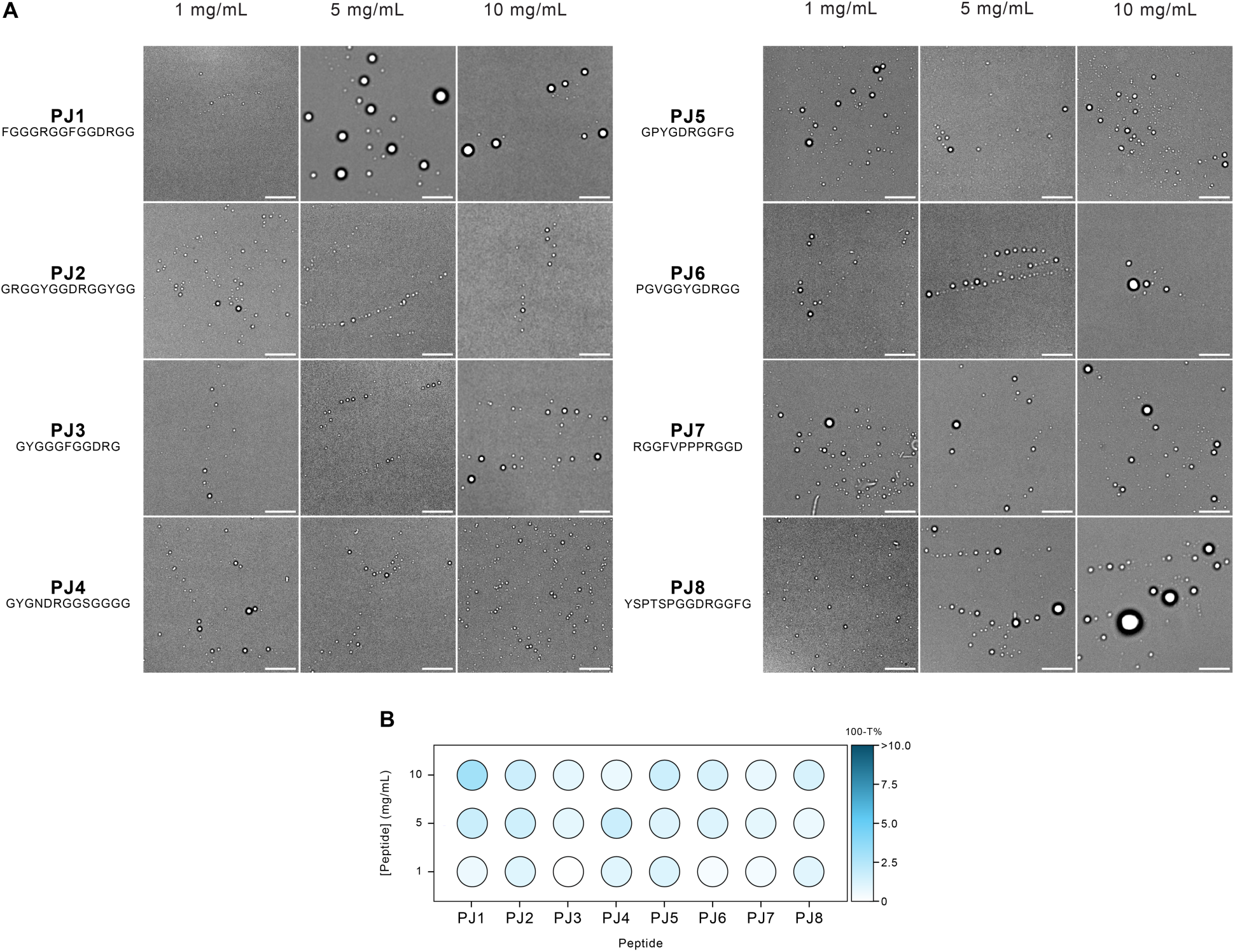
(A) Bright-field microscope images depicting peptide-induced LLPS at various concentrations. Scale bar: 10 μm. (B) Phase diagrams illustrating the effect of different peptide concentrations on the LLPS of PJ peptides, measured by the relative turbidity (100-T%). Different blue shades represent different relative turbidity levels.

Our observations revealed that all samples produced droplets across the full range of peptides and concentrations tested. In our turbidity measurements, we observed a general trend of increased turbidity at higher concentrations across all samples. Notably, PJ1 had one of the most effective droplet formation and the highest turbidity.

PJ1, our highest-scoring designed peptide, exhibited enhanced condensate formation at a concentration of 5 mg/mL, with notable abundance and size of the liquid droplets (2-5μm), in line with previous studies in literature. ^11,13,14,17^ This result is particularly significant as it demonstrates that our peptide design method’s top score indeed corresponded to the best LLPS performance.

The PJ8 peptide, containing a partial sequence of the established LLPS-associated YSPTSPSY motif, exhibited enhanced droplet formation as well. Notably, PJ8 was not selected primarily for its high score in our design method; its LLPS behavior likely originates from a mechanism distinct from PJ1’s co-occurrence of motif trios. This observation suggests that even a partial LLPS motif alone can promote phase separation, independent of other motifs within the short sequence. This finding shows an alternative mechanism for LLPS in designed peptides, presenting a promising design strategy for future research.

Apart from PJ1 and PJ8, varying the peptide scores does not significantly impact phase separation, as all our sequences contain LLPS-prone motifs. Consequently, the LLPS propensity of the designed peptides is indistinguishable, with all of them undergoing phase separation.

Fluorescence Recovery After Photobleaching (FRAP) was implemented as a powerful technique to study the dynamics and liquid-like character of the peptide condensates, by partitioning the FITC guest molecule and performing fluorescence recovery of bleached droplets over time. To ensure comparability across all PJ peptides, we maintained consistent FRAP parameters (i.e. laser intensity, gain settings, etc.) for all experiments.

The FRAP experiment results for the PJ peptide sequences provide evidence for their liquid-like character, as all peptides exhibited recovery percentages ranging from 36% (PJ4) to 92% (PJ5), as shown in Fig. 7. This overall recovery across the peptide set confirms their intrinsic ability for molecular diffusion and local rearrangement within their interior. ^25,57,77^ The wide range of recovery percentages observed suggests that while all peptides display liquid-like behavior, their specific sequence compositions significantly influence their mobility and diffusion dynamics.

**Figure 7.**
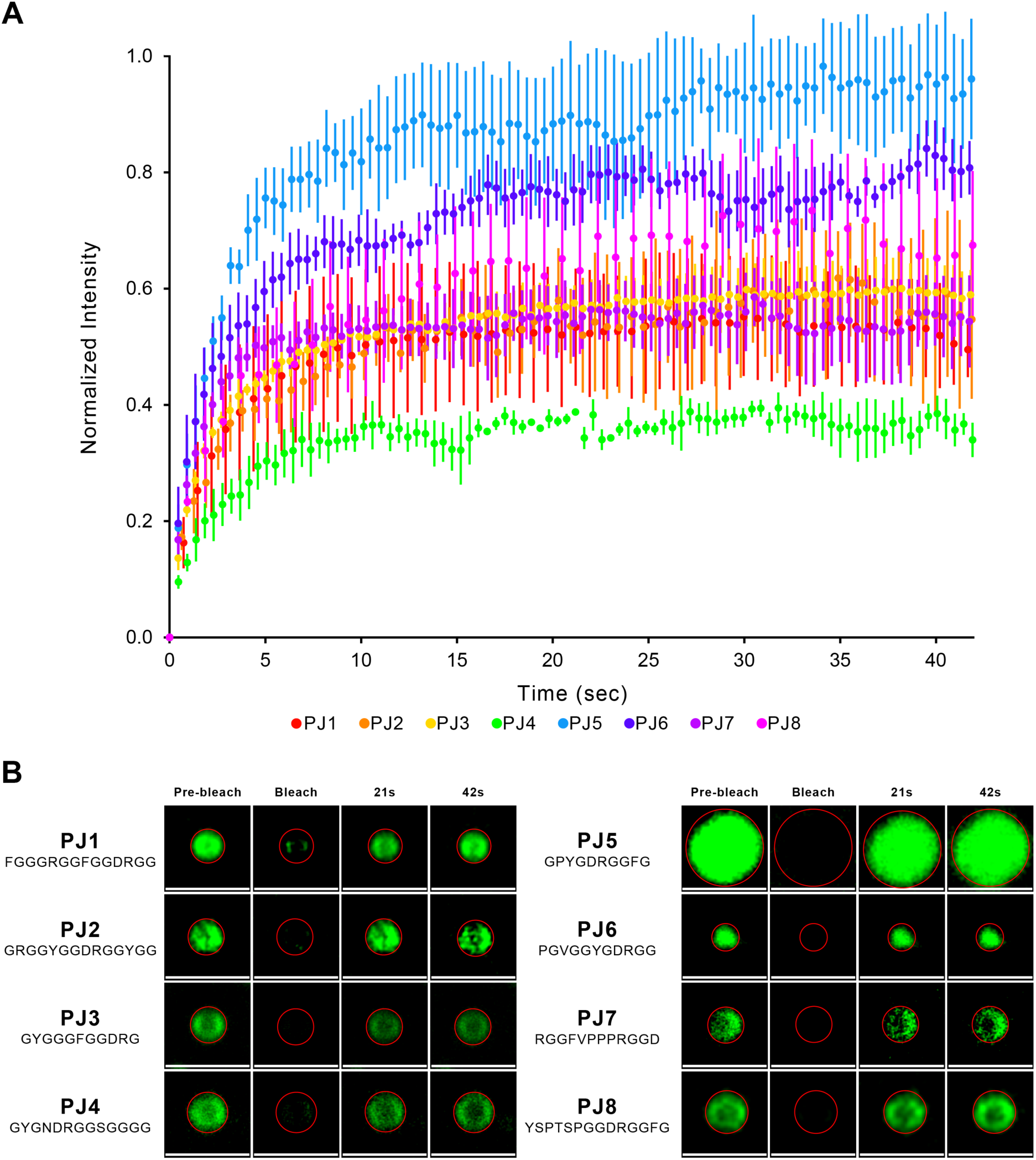
Ability of designed peptides to undergo LLPS and form liquid-like droplets. (A) FRAP analysis of peptide condensates (data is presented as mean values ± SD of 3 condensates), (B) Representative confocal images illustrating the FRAP process in an individual condensate at different time points. Scale bars set to 2 μm.

While PJ1, the peptide with the highest score of our peptide design based on motif co-occurrence, and PJ8, which includes a partial sequence of the established LLPS-associated YSPTSPSY, exhibited the most effective droplet formation previously (Fig. 6), and the FRAP experiments confirmed their liquid-like characteristics, they did not exhibit the highest recovery percentages among the peptide set. This suggests that our scoring system and the presence of LLPS motifs does not correlate with the dynamics of condensate behavior. Instead, it appears that the dynamics are influenced by other factors, including specific sequence composition.

Among the analyzed peptides, PJ5, the shortest at 10 residues, exhibited the highest recovery percentage at 92%. Its sequence composition and characteristics reflect a balanced combination of features from the entire PJ peptide set, including Gly and Pro relative content, as well as polar and hydrophobic properties - key factors that are known to contribute to LLPS. ^3,4,7^

By categorizing the PJ set into peptides with the highest scores in our peptide design method (PJ1-4, scores 55-74) and those with the lowest scores (PJ5-8, scores 15-29), as detailed in Table 2, we can identify notable trends. Peptides PJ1-4 exhibit recovery percentages of 36% to 58%, while PJ5-8 show higher recovery rates at 54% to 92%.

The lower recoveries of PJ1-4 peptides, which were designed using motifs that frequently co-occur in DPR sequences, suggest that these motifs may engage in inter-/intra-molecular interactions, which benefit the formation of a more stable and less flexible network within the droplets. ^60,78,79^ Such stability could reduce the fluidity of the droplets, thereby limiting the exchange of molecules between the droplets and their environment. ^80^

In contrast, the droplets formed by PJ5-8 peptides exhibit enhanced liquid-like characteristics, as confirmed by their higher fluorescence recoveries. These fluid structures are characterized by rapid organization, facilitating the easy entry, diffusion, and exit of macromolecules, ^17,29,77,81^ which can be attributed to reduced constraints from interactions, as low-scoring peptides contain motifs that do not frequently co-occur within DPRs. ^29,60,79,80,82^ As a result, fluorescence in laser-bleached regions within these droplets can recover quickly due to this more efficient molecular exchange.^29,60,80,81^

Interestingly, PJ5-8 are the only Pro-containing peptides, which could be another factor that explains the improved fluorescence recoveries. Pro, with its unique propensity to form a cyclic structure, is known to disrupt secondary structures in proteins. ^51,83^ However, in small peptides, Pro can induce folding rather than disrupt it by creating a distinctive kink that can promote turns or bends. ^11,51,84–88^ This structural change can influence droplet dynamics by promoting peptide folding, which has been shown to enhance the liquid-like character of condensates. ^17,89^ This increased fluidity would lead to higher fluorescence recoveries in Pro-containing peptides. With this, further research regarding PJ secondary structures is needed to confirm such hypotheses.

The experimental data validates our computational peptide-design strategy, with all PJ peptides demonstrating LLPS behavior across various concentrations. The FRAP experiments further shows the liquid-like nature of our condensates, revealing complex relationships between sequence features and droplet dynamics.

## Conclusions

This study presents a novel approach for identifying and characterizing liquid-liquid phase separation (LLPS) short motifs across diverse Phase-Separating Proteins. Our analysis revealed a complex landscape of LLPS-associated sequences, encompassing both known and previously unidentified motifs. The discovery of family-specific motif variations highlights the intricate relationship between protein function and phase separation propensity.

We developed a non-biased computational method for minimalistic peptide design based on the co-occurrence of the discovered motifs in Droplet Promoting Regions of Phase-Separating Proteins. By selecting a diverse array of peptides generated through this method, we confirmed that all of them undergo LLPS, albeit potentially through different mechanisms. Notably, the different primary sequences resulted in distinct liquid droplet dynamics and behaviors, underscoring the significant influence of sequence composition on phase separation characteristics.

Our research bridges the gap between primary structure and phase separation properties, revealing that sequence determinants of LLPS are more diverse and nuanced than previously recognized. These insights provide a foundation for future work in biomolecular engineering, potentially enabling the manipulation of protein and peptide sequences to design biocondensates with tailor-made properties. Our work sets a precedent for creating functional biomolecular assemblies, with significant implications for therapeutic interventions in phase separation-related disorders and the development of engineered peptide-based materials.

## Code availability

The code underlying this study are openly available in our repository *UnderstandingLLPS* at https://github.com/BPSlab/UnderstandingLLPS.git.

## Acknowledgements

The authors thank Prof. Monika Fuxreiter from the University of Padova and Prof. Leonor Morgado and Prof. Cláudio Soares from ITQB NOVA for the helpful discussions, as well as Dr. Carolina Feliciano and Dr. Pedro Matos Pereira from Bacterial Imaging Cluster (BIC) at ITQB NOVA for the help with the microscopy experiments.

This work was supported by FCT - Fundação para a Ciência e a Tecnologia, I.P., through MOSTMICRO-ITQB R&D Unit (DOI 10.54499/UIDB/04612/2020; DOI 10.54499/UIDP/04612/2020;), LS4FUTURE Associated Laboratory (DOI 10.54499/LA/P/0087/2020), UI/BD/154577/2022 for J.C. and 2021.01283.CEECIND/CP1657/CT0004 (DOI 10.54499/2021.01283.CEECIND/CP1657/CT0004) for A.S.P. This work was partially supported by PPBI - Portuguese Platform of BioImaging (PPBI-POCI-01-0145-FEDER-022122) co-funded by national funds from OE - “Orçamento de Estado” and by european funds from FEDER - “Fundo Europeu de Desenvolvimento Regional”.

The authors would also like to thank the Croatian Science Foundation/Hrvatska zaklada za znanost (grant no: UIP-2019-04-7999) and the University of Rijeka for support.

