## Supplementary Information for "Investigating Amino acid Enrichments and Patterns in Phase-Separating Proteins: Understanding Biases in Liquid-Liquid Phase Separation"

### 1. Supplementary Notes

#### 1.1 Family-specific motif variations

In the general motif discovery phase, we identified 129 motifs with a CF ≥ 0.2. For the protein families, CF thresholds were adjusted to yield comparable motif counts: RNA binding (CF ≥ 0.35, 90 motifs), DNA binding (CF ≥ 0.30, 72 motifs), Chromatin binding (CF ≥ 0.40, 66 motifs), Regulation (CF ≥ 0.25, 74 motifs), Hydrolases (CF ≥ 0.40, 84 motifs), and Structure (CF ≥ 0.30, n=107).

#### 2. Supplementary Figures

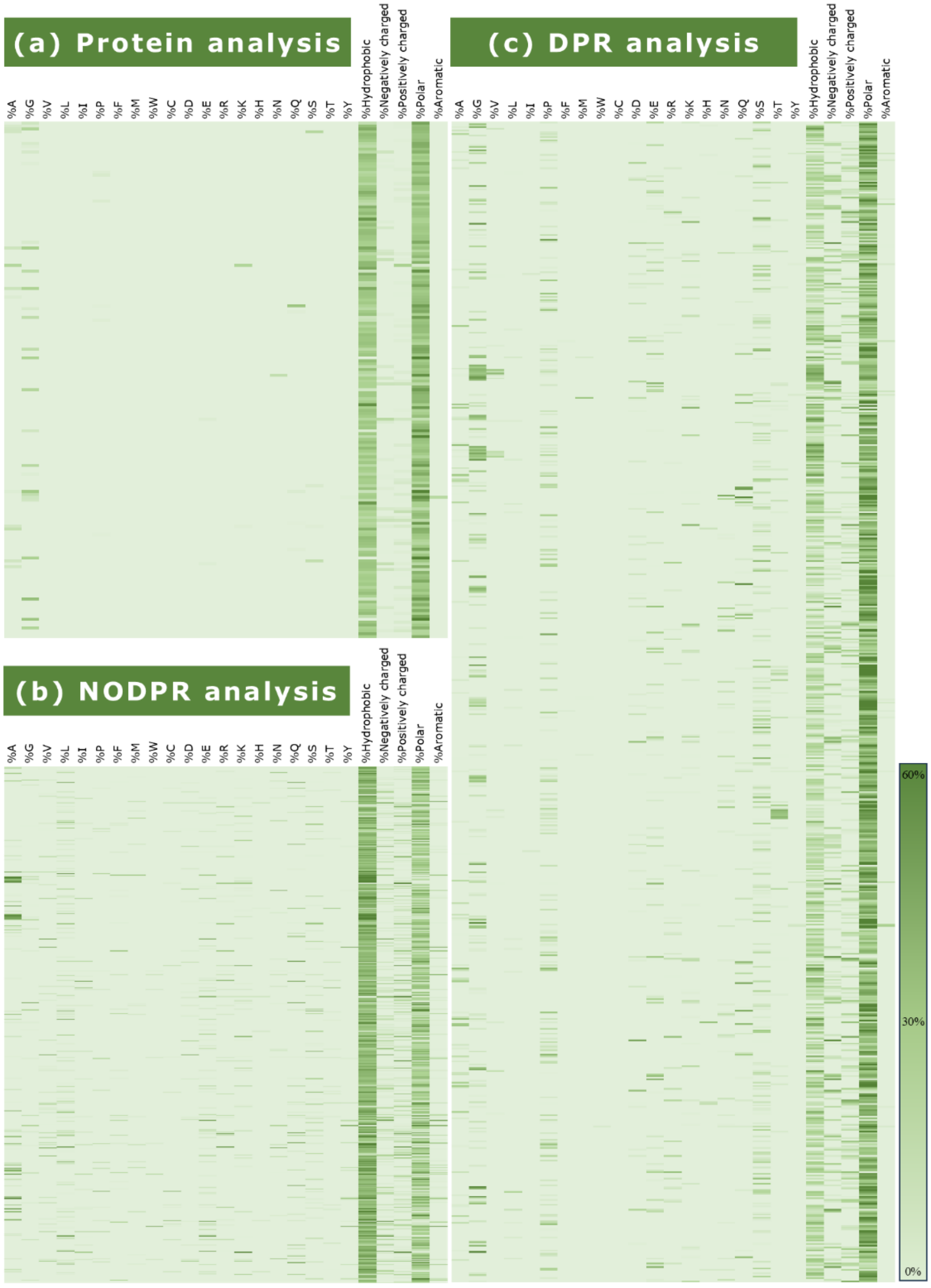


*Figure S1 - Analysis of amino acids and residue character enrichment in (a) PhSePs; (b) NODPRs; (c) DPRs. The color gradient transitions from lighter hues (corresponding to 0%) to darker shades (indicating the maximum value observed at 60% presence).*

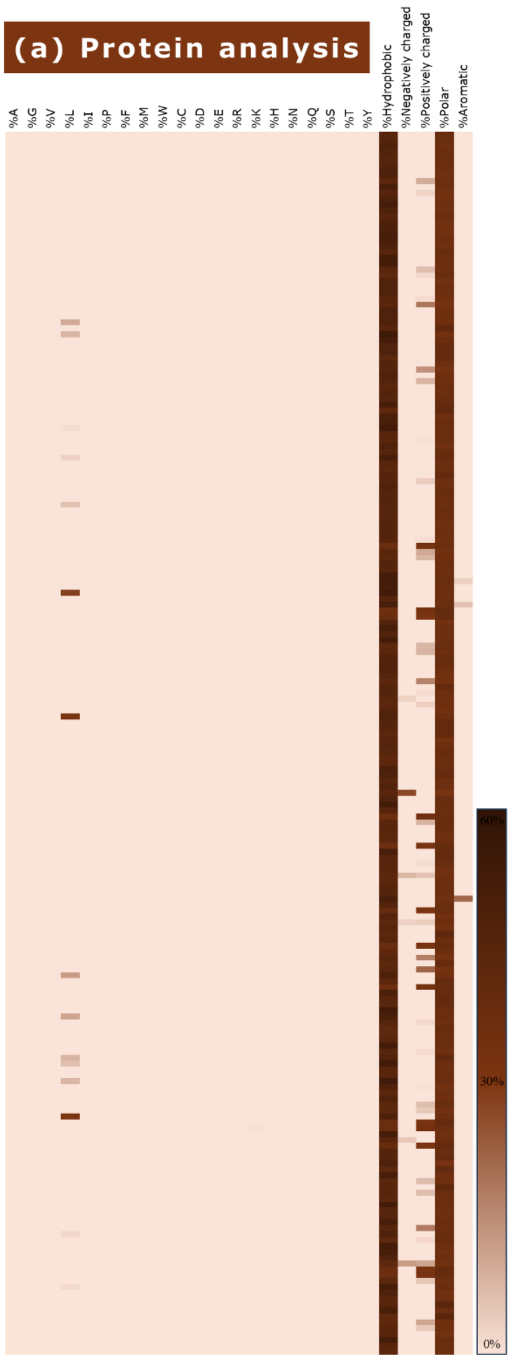


*Figure S2 - Analysis of amino acids and residue character enrichment in (a) non-PhSePs. The color gradient transitions from lighter hues (corresponding to 0%) to darker shades (indicating the maximum value observed at 60% presence).*

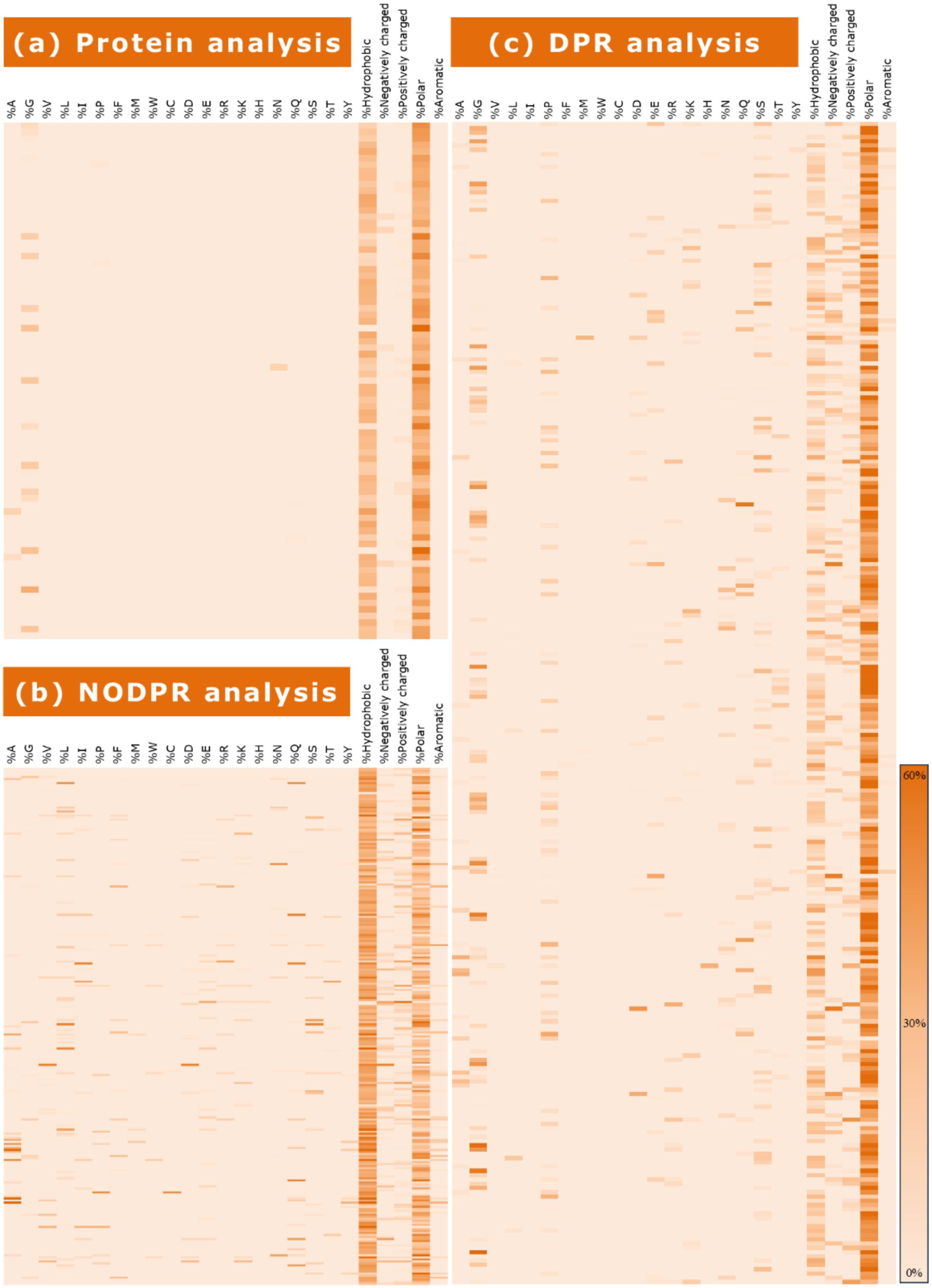


*Figure S3 - Analysis of amino acids and residue character enrichment in the RNA binding family, (a) PhSePs; (b) NODPRs; (c) DPRs. The color gradient transitions from lighter hues (corresponding to 0%) to darker shades (indicating the maximum value observed at 60% presence).*

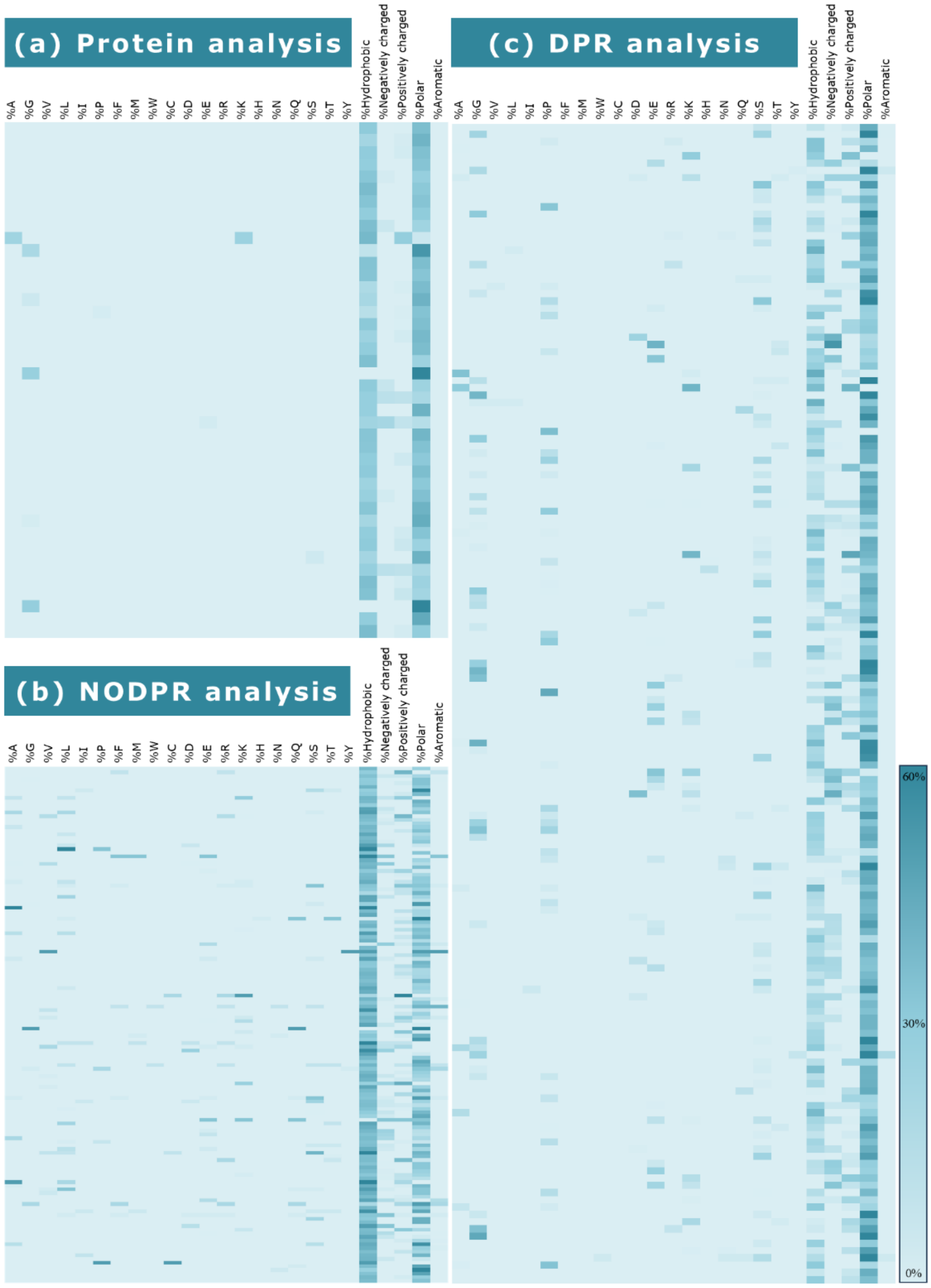


*Figure S4 - Analysis of amino acids and residue character enrichment in the DNA binding family, (a) PhSePs; (b) NODPRs; (c) DPRs. The color gradient transitions from lighter hues (corresponding to 0%) to darker shades (indicating the maximum value observed at 60% presence).*

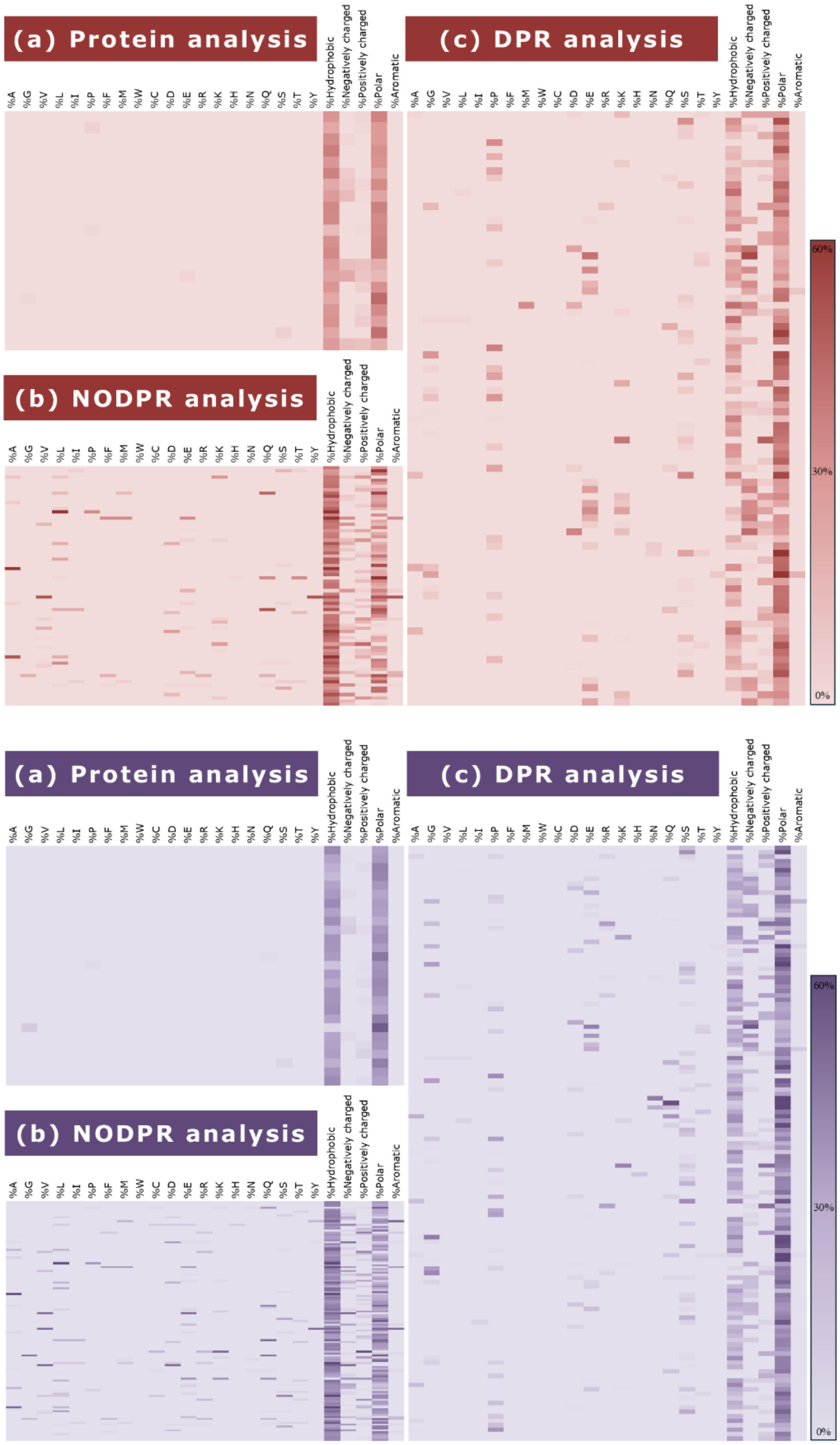


*Figure S5 - Analysis of amino acids and residue character enrichment in the Chromatin binding (red) and Regulation (purple) families, (a) PhSePs; (b) NODPRs; (c) DPRs. The color gradient transitions from lighter hues (corresponding to 0%) to darker shades (indicating the maximum value observed at 60% presence).*

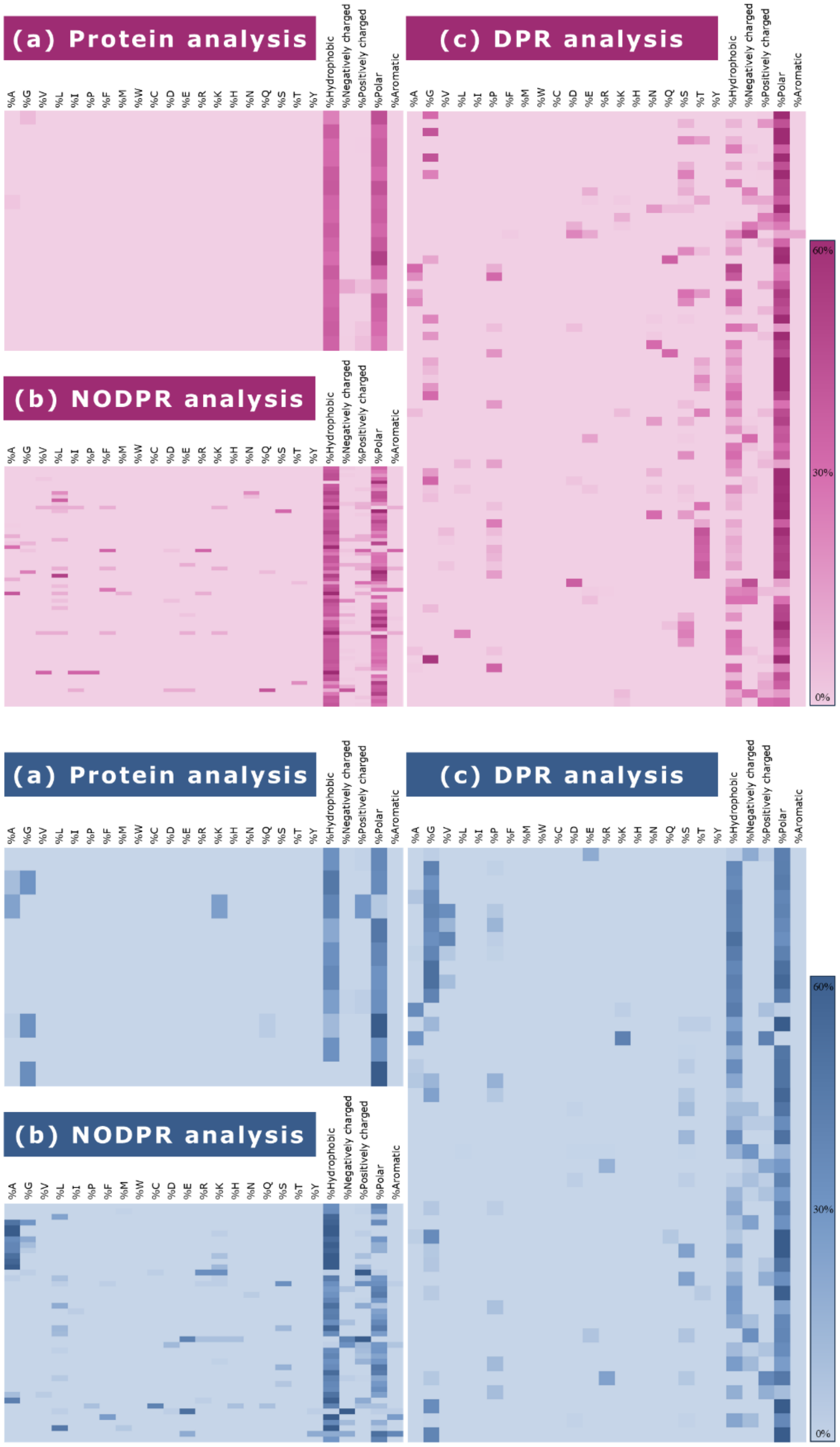


*Figure S6 - Analysis of amino acids and residue character enrichment in the Hydrolase (pink) and Structure (dark blue) families, (a) PhSePs; (b) NODPRs; (c) DPRs. The color gradient transitions from lighter hues (corresponding to 0%) to darker shades (indicating the maximum value observed at 60% presence).*

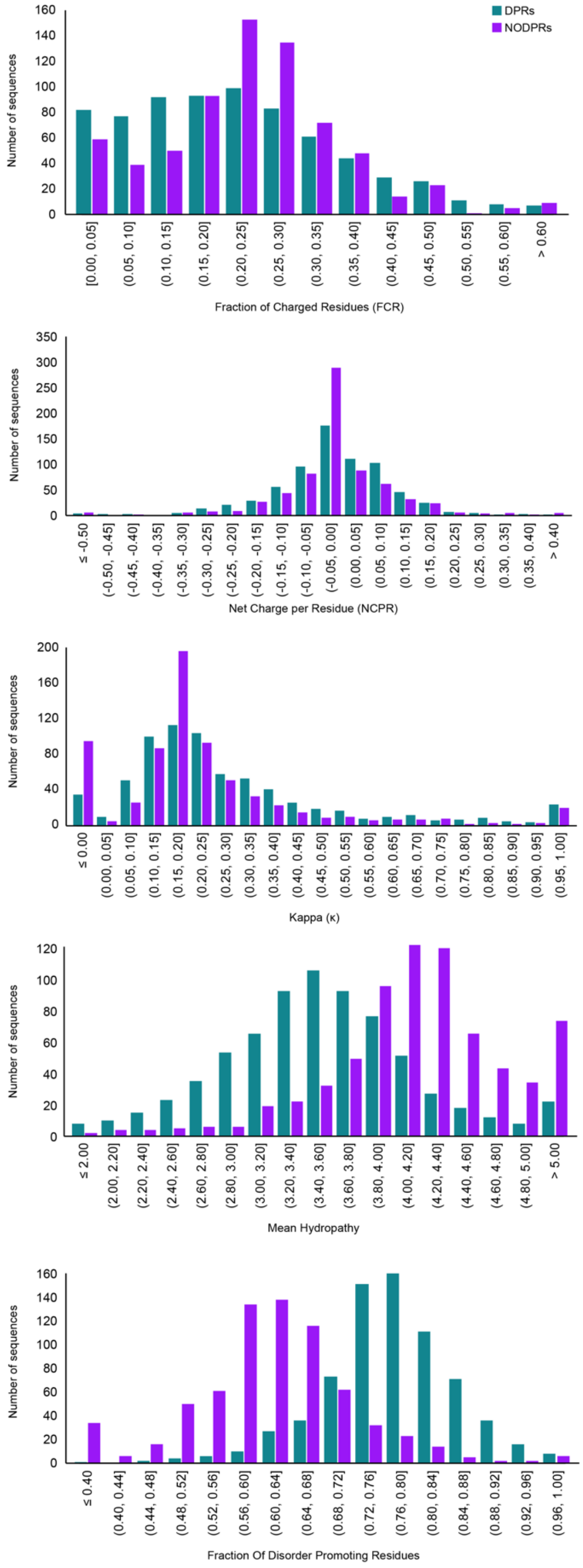


*Figure S7 – Cider Server parameter distribution for DPRs and NODPRs.*

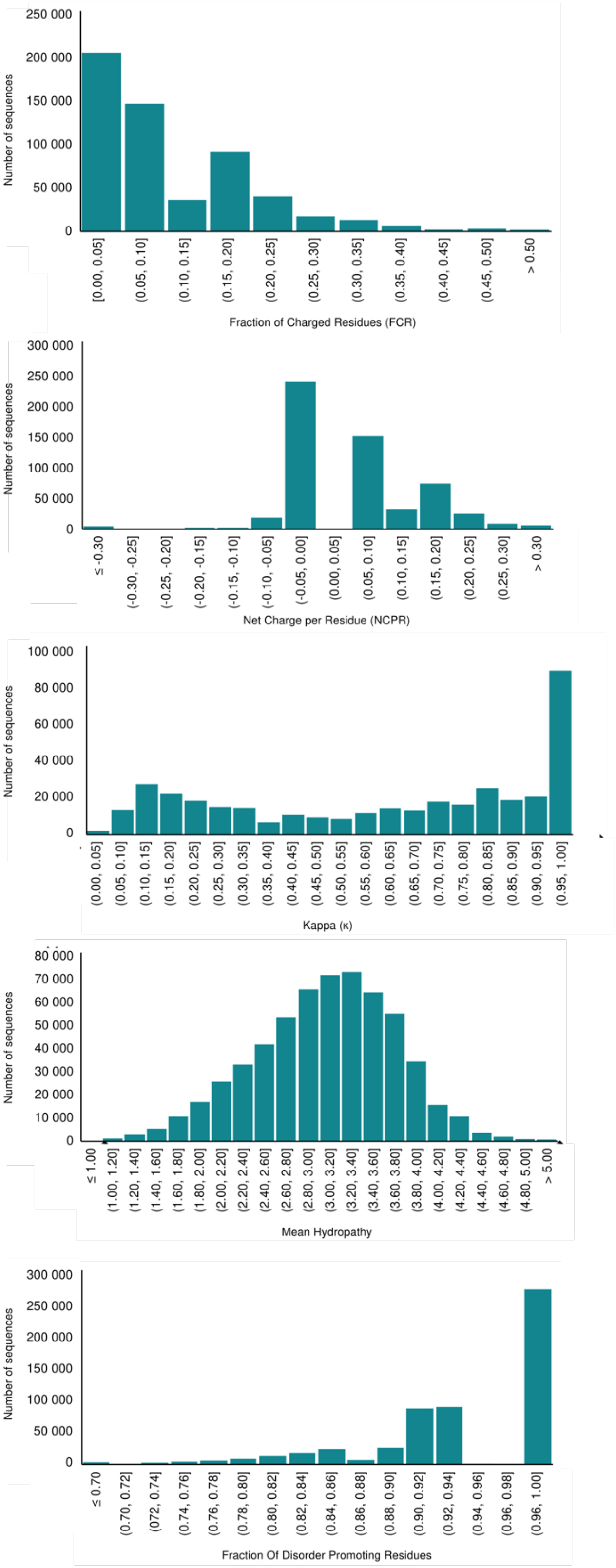


*Figure S8 - Cider Server parameter distribution for designed peptides.*

#### 3. Supplementary Tables

*Table S1 – Presence and Frequency values of discovered motifs in DPRs and NODPRs.*

| Motif | NODPR | | DPR | |
|---|---|---|---|---|
| | Presence | Frequency | Presence | Frequency |
| AAPA | 1 | 1 | 13 | 23 |
| DDED | 1 | 1 | 11 | 12 |
| DEDD | 1 | 1 | 11 | 13 |
| DRGG | 0 | 0 | 13 | 39 |
| DSSS | 2 | 2 | 11 | 21 |
| FGGG | 1 | 1 | 13 | 20 |
| GAPG | 0 | 0 | 11 | 27 |
| GDRG | 0 | 0 | 14 | 39 |
| GDRGG | 0 | 0 | 11 | 32 |
| GFGG | 1 | 1 | 18 | 28 |
| GGDR | 2 | 2 | 13 | 44 |
| GGDRG | 0 | 0 | 12 | 37 |
| GGDRGG | 0 | 0 | 11 | 32 |
| GGFGG | 0 | 0 | 10 | 16 |
| GGGG | 1 | 1 | 40 | 98 |
| GGGGF | 0 | 0 | 10 | 13 |
| GGGGG | 1 | 1 | 25 | 51 |
| GGGGGG | 0 | 0 | 14 | 25 |
| GGGGR | 0 | 0 | 11 | 16 |
| GGGGS | 0 | 0 | 13 | 16 |
| GGGN | 2 | 2 | 14 | 29 |
| GGGR | 2 | 2 | 26 | 35 |
| GGGRG | 0 | 0 | 17 | 25 |
| GGGRGG | 0 | 0 | 14 | 16 |
| GGGSG | 0 | 0 | 15 | 20 |
| GGGSGG | 0 | 0 | 11 | 13 |
| GGGY | 0 | 0 | 15 | 32 |
| GGGYG | 0 | 0 | 13 | 22 |
| GGGYGG | 0 | 0 | 12 | 20 |
| GGNG | 0 | 0 | 10 | 19 |
| GGPGG | 0 | 0 | 10 | 13 |
| GGPP | 0 | 0 | 12 | 13 |
| GGRG | 1 | 1 | 29 | 70 |
| GGRGG | 0 | 0 | 22 | 46 |
| GGRS | 0 | 0 | 11 | 11 |
| GGSGG | 0 | 0 | 15 | 21 |
| GGSS | 2 | 2 | 12 | 15 |
| GGYG | 1 | 1 | 23 | 74 |
| GGYGG | 0 | 0 | 17 | 47 |
| GNGG | 1 | 1 | 8 | 19 |
| GPGS | 1 | 1 | 9 | 21 |
| GPPP | 0 | 0 | 16 | 22 |
| GPYG | 0 | 0 | 9 | 21 |
| GRGG | 0 | 0 | 34 | 86 |
| GRGGG | 0 | 0 | 15 | 25 |
| GRGGY | 0 | 0 | 10 | 14 |
| GRGR | 1 | 1 | 13 | 15 |
| GRGS | 0 | 0 | 10 | 13 |
| GSGGG | 0 | 0 | 10 | 14 |
| GVPGV | 0 | 0 | 8 | 19 |
| GYGGG | 0 | 0 | 9 | 16 |
| GYGN | 1 | 1 | 11 | 11 |
| HHP | 2 | 2 | 13 | 16 |
| HQQQ | 0 | 0 | 15 | 21 |
| HQQQQ | 0 | 0 | 12 | 16 |
| PAPA | 1 | 1 | 17 | 31 |
| PGGG | 1 | 4 | 16 | 25 |
| PGGP | 0 | 0 | 14 | 17 |
| PGQQ | 1 | 1 | 5 | 35 |
| PGVG | 1 | 2 | 18 | 43 |
| PGVGV | 1 | 1 | 8 | 23 |
| PPPG | 2 | 2 | 22 | 25 |
| PPPP | 4 | 4 | 38 | 70 |
| PPPPP | 0 | 0 | 19 | 35 |
| PPPPPP | 0 | 0 | 10 | 16 |

*Table S1 - continued*

| Motif | NODPR | | DPR | |
|---|---|---|---|---|
| | Presence | Frequency | Presence | Frequency |
| PPPPQ | 0 | 0 | 11 | 14 |
| PPPQ | 0 | 0 | 22 | 35 |
| PPQG | 1 | 1 | 12 | 13 |
| PPSS | 0 | 0 | 14 | 15 |
| PQQP | 1 | 1 | 11 | 16 |
| PQQQ | 0 | 0 | 19 | 32 |
| PQQQQ | 0 | 0 | 14 | 20 |
| PSGP | 0 | 0 | 12 | 14 |
| PSSS | 2 | 2 | 12 | 17 |
| PSYS | 0 | 0 | 2 | 46 |
| PSYSP | 0 | 0 | 2 | 46 |
| PSYSPT | 0 | 0 | 2 | 42 |
| PTSPSY | 0 | 0 | 2 | 42 |
| QGPG | 1 | 1 | 4 | 46 |
| QPN | 3 | 3 | 15 | 18 |
| QPPPP | 1 | 1 | 12 | 18 |
| QPQQ | 1 | 1 | 12 | 18 |
| QQGP | 0 | 0 | 2 | 45 |
| QQPP | 2 | 2 | 17 | 28 |
| QQPPP | 1 | 1 | 11 | 18 |
| QQPQ | 0 | 0 | 13 | 18 |
| QQQ | 18 | 19 | 58 | 199 |
| QQQH | 1 | 1 | 10 | 18 |
| QQQP | 1 | 1 | 24 | 31 |
| QQQQ | 6 | 6 | 36 | 110 |
| QQQQP | 0 | 0 | 13 | 17 |
| QQQQQ | 4 | 4 | 28 | 64 |
| QQQQQP | 0 | 0 | 11 | 12 |
| QQQQQQ | 1 | 1 | 22 | 46 |
| RGGD | 0 | 0 | 12 | 16 |
| RGGF | 1 | 1 | 19 | 29 |
| RGGFG | 0 | 0 | 10 | 13 |
| RGGG | 1 | 1 | 26 | 50 |
| RGGGG | 0 | 0 | 14 | 24 |
| RGGR | 2 | 2 | 23 | 43 |
| RGGRG | 0 | 0 | 15 | 34 |
| RGGRGG | 0 | 0 | 11 | 24 |
| RGRG | 1 | 1 | 24 | 32 |
| RGRGG | 0 | 0 | 17 | 21 |
| RRGG | 1 | 1 | 13 | 16 |
| SAPA | 0 | 0 | 12 | 23 |
| SGGGG | 0 | 0 | 13 | 19 |
| SGGGGG | 0 | 0 | 11 | 15 |
| SPSYS | 0 | 0 | 2 | 46 |
| SPSYSP | 0 | 0 | 2 | 46 |
| SPTSP | 0 | 0 | 4 | 63 |
| SPTSPS | 0 | 0 | 3 | 41 |
| SQQP | 0 | 0 | 12 | 12 |
| SRGG | 1 | 1 | 19 | 20 |
| SSAP | 2 | 2 | 10 | 22 |
| SSDS | 1 | 1 | 13 | 16 |
| SSSD | 6 | 6 | 11 | 16 |
| SSTG | 2 | 2 | 11 | 14 |
| SYSPT | 0 | 0 | 2 | 42 |
| SYSPTS | 0 | 0 | 2 | 42 |
| TSPSY | 0 | 0 | 2 | 42 |
| TSPSYS | 0 | 0 | 2 | 42 |
| VPGVG | 0 | 0 | 9 | 22 |
| VPPP | 3 | 3 | 17 | 17 |
| YGGG | 0 | 0 | 14 | 30 |
| YGPG | 1 | 1 | 6 | 29 |
| YSPT | 0 | 0 | 2 | 61 |
| YSPTS | 0 | 0 | 2 | 60 |
| YSPTSP | 0 | 0 | 2 | 60 |

*Table S2 – Unique motifs identified for each protein family.*

| RNA binding | DNA binding | Chromatin binding | Regulation | Hydrolase | | Structure | |
|---|---|---|---|---|---|---|---|
| FRGGRG | GGSGG | ERRRRE | AQAQ | AASSS | NVGDTT | AAAK | GVGG |
| GDRGGF | GGSGGG | HHQ | AQAQA | ATPT | PATPT | AAAKK | GVGGL |
| GGFGG | GRGRG | KSKK | ASSPG | ATPTT | PATPTT | AAAKKK | GVGGLG |
| GGFGGG | GSGGG | MGQ | GGPGG | ATPTTP | PPHLR | AAAKKP | GVKP |
| GGFRGG | GSGGGG | NTWE | GGPGGP | DEDD | PTTP | AAGG | GVLPG |
| GGGGGF | GYRGRG | NTWEP | GPGGP | DRGGR | PTTPV | AAKK | GVLPGV |
| GGGGR | HHT | PKH | HPSSM | DRGGRG | PTTPVT | AAKKP | GVPG |
| GGR | PSYS | PPGA | PAPA | DRGR | PVTTP | AAKKPK | GVPGV |
| GGRGGD | PSYSP | PPP | PAPAP | DTT | TEPA | AAKP | GVPGVG |
| GGRGGY | PSYSPT | PPPG | PASSS | DTTE | TEPAT | AGG | KAAAKK |
| GGYG | PTSPSY | PPPH | PPASS | DTTEP | TEPATP | AGGY | KKDG |
| GGYGGD | RGRGGG | PPPLP | PPSS | DTTEPA | TGFG | AKKPKK | KKPKKA |
| GGYGGG | SPSYS | PPPPPP | PPTPP | EPATP | TGTA | APGAIP | KPA |
| GRG | SPSYSP | PPPPPQ | PPTT | EPATPT | TPTT | DENK | KPKA |
| GRGGDR | SPT | PPPPQ | PQQQ | GDRGGR | TPTTP | ETSK | KPKKA |
| GRGGF | SPTSP | PPPQQ | PQQQQ | GDTT | TPTTPV | GAGG | LGVGG |
| GRGGY | SPTSPS | PQM | PTPP | GDTTE | TPVTT | GAGGAG | LGVGGL |
| GYGGDR | SRGGG | PQQPPP | QAQA | GDTTEP | TPVTTP | GAKA | LPGVG |
| GYGGG | SRGGGG | QHH | QAQAQ | GGGGF | TTEP | GAPG | LPGVGV |
| HQQQQQ | SSFSSS | QPPPPP | QAQAQA | GGGGY | TTEPA | GAPGA | LSFG |
| MGG | SSSFSS | QQQPP | QQPQ | GGLFG | TTEPAT | GAPGAI | NGNG |
| MGM | SSSSST | REQ | QQPS | GGVNV | TTPV | GAVPG | NGNGG |
| NDN | SYSPT | SSDS | QRSA | GGVNVG | TTPVT | GFGPGG | PGAIPG |
| NYN | SYSPTS | SSSD | RGGPG | GLFG | TTPVTT | GGGN | PGAPG |
| PPM | TSPSY | SSSDS | RGGPGG | GVNVG | VGDTT | GGLG | PGAPGA |
| PPQ | TSPSYS | TPP | RGRGR | GVNVGD | VGDTTE | GGLGV | PGAVPG |
| RGGDRG | YRGRGG | TWE | RQQQQ | MYS | VNVGD | GGLGVG | PGG |
| RGGFG | YSPT | TWEP | SAAAA | NGGGG | VNVGDT | GGNG | PGGQ |
| RGGFGG | YSPTS | TWEPE | SHSR | NGGGGG | VTTP | GGNGN | PGGR |
| RGGGG | YSPTSP | WEPE | SPSSD | NVGD | YGGDRG | GGNGNG | PGGRP |
| RGGGGG | | | SQGEE | NVGDT | | GGRP | PGGV |
| RGGYG | | | SQPQ | | | GGVGGL | PGGVAG |
| RGGYGG | | | SSHS | | | GGVPG | PGIG |
| YGG | | | SSTG | | | GLGV | PGLG |
| YGGG | | | TPPTT | | | GLGVG | PGQ |
| YND | | | TPTA | | | GLGVGG | PGQG |
| YQA | | | | | | GLPG | PGVG |
| | | | | | | GLPGV | PGVGV |
| | | | | | | GLPGVG | PGVYPG |
| | | | | | | GNG | PKKA |
| | | | | | | GNGG | QQG |
| | | | | | | GNGN | QSD |
| | | | | | | GNGNG | SGGR |
| | | | | | | GNGNGG | SGGRP |
| | | | | | | GPGG | SGPGG |
| | | | | | | GPGGQ | VGAGV |
| | | | | | | GPY | VGGLG |
| | | | | | | GPYG | VGGLGV |
| | | | | | | GQGG | VKPK |
| | | | | | | GRDG | VPGA |
| | | | | | | GSGP | VPGVG |
| | | | | | | GVGAG | VPGVGV |
| | | | | | | GVGAGV | YGAP |
